# Memory sequencing reveals heritable single cell gene expression programs associated with distinct cellular behaviors

**DOI:** 10.1101/379016

**Authors:** Sydney M. Shaffer, Benjamin L. Emert, Raul Reyes-Hueros, Christopher Coté, Guillaume Harmange, Ann E. Sizemore, Rohit Gupte, Eduardo Torre, Abhyudai Singh, Danielle S. Bassett, Arjun Raj

## Abstract

Non-genetic factors can cause individual cells to fluctuate substantially in gene expression levels over time. Yet it remains unclear whether these fluctuations can persist for much longer than the time of one cell division. Current methods for measuring gene expression in single cells mostly rely on single time point measurements, making the duration of gene expression fluctuations or *cellular memory* difficult to measure. Here, we report a method combining Luria and Delbrück’s fluctuation analysis with population-based RNA sequencing (MemorySeq) for identifying genes transcriptome-wide whose fluctuations persist for several cell divisions. MemorySeq revealed multiple gene modules that are expressed together in rare cells within otherwise homogeneous clonal populations. Further, we found that these rare cell subpopulations are associated with biologically distinct behaviors, such as the ability to proliferate in the face of anti-cancer therapeutics, in different cancer cell lines. The identification of non-genetic, multigenerational fluctuations has the potential to reveal new forms of biological memory at the level of single cells and suggests that non-genetic heritability of cellular state may be a quantitative property.

## Main text

Cellular memory in biology, meaning the persistence of a cellular or organismal state over time, occurs over a wide range of timescales and can be produced by a variety of mechanisms. Genetic differences are one form of memory (Ben-David et al., 2018), encoding variation between organisms on multi-generational timescales. Within an organism, mechanisms involving the regulation of gene expression encode the differences between cell types in different tissues, with cells retaining memory of their state over a large number of cell divisions (Bonasio et al., 2010). In contrast, recent measurements suggest that the expression of many genes in single cells may have very little memory, displaying highly transient fluctuations in transcription. These rapid fluctuations have been referred to as gene expression “noise” and have generally been difficult to associate with physiological distinctions between single cells (Raj and van Oudenaarden, 2008; Raj et al., 2006; Sigal et al., 2006; Symmons and Raj, 2016), although there are certainly specific examples in which such fluctuations can drive phenotype (Cohen et al., 2008; Raj et al., 2010; Wernet et al., 2006).

Less well studied is memory on intermediate timescales; i.e., cellular states that are ultimately transient, and thus are not indefinitely heritable, but may persist for several divisions, and thus are not “noise” either. Such timescales would be long enough to allow for coordinated fluctuations in the expression of many genes at once in individual cells, potentially resulting in biological activity within that cell that is distinct from the rest of the population. Yet it remains unclear how prevalent such longer duration fluctuations might be, because finding the molecular markers of these longer fluctuations is difficult: current “snapshot” methods are unable to distinguish between fast and slow fluctuations because they lack any temporal component, while time lapse microscopy is laborious and difficult to scale to all genes (Hormoz et al., 2016; Phillips et al., 2019). Thus, we sought to develop a method that would enable us to find genes whose expression fluctuations would be maintained over several cell divisions. Ultimately, our goal was to use these markers of slow fluctuations to identify functionally distinct subpopulations within otherwise indistinguishable cells.

The methodology we developed to distinguish heritable from non-heritable fluctuations in expression levels in single cells (MemorySeq) is based on the fluctuation analysis from Luria and Delbrück’s beautiful 1943 experiments on resistance to phage in bacteria, which they used to discriminate heritable from non-heritable mechanisms for resistance (Luria and Delbrück, 1943) (also used in cancer (Shaffer et al., 2017; Tlsty et al., 1989)). In our context of cellular memory, the experiment consisted of growing a number of “MemorySeq clones” (we aimed for 48 and ended up with 42-45 after losses from culture and library prep; see Methods for details) of isogenic melanoma cells (WM989-A6) in individual wells, eventually growing them to around 100,000 cells per clone (Fig. 1A,B). If a fluctuating gene transitioned in and out of the “high” expression state relative rapidly compared to the cell division rate, then a fairly constant proportion of those 100,000 cells would be in the high expression state for that gene (with some dispersion due to Poisson sampling). This constancy occurs because the cells do not remember the state through cell division. At the opposite extreme, if the high expression state was long-lived compared to the cell division time, then if a cell occasionally moves into the high expression state early in the family tree, all of its progeny will remain in the high expression state, leading to a very high proportion of the final 100,000 cells being in the high expression state. Thus, across multiple MemorySeq clones, we would find a high variance in the proportion of cells in the high expression state in the final population, where most clones would have low expression of that gene and a few clones would have high expression, depending on exactly how far up in the family trees the cells transitioned into the high expression state. To measure variability in the proportion of cells in the high expression state for any particular gene, we used bulk RNA sequencing to measure the transcription of all genes in each expanded clone, and we then measured the variability in the expression of all genes across these clones (Fig. 1B). It was important to distinguish between biological variability between clones and variability due to sequencing, sampling, and technical errors. We therefore also grew a large population of cells that we split into 48 wells containing 100,000 cells each and subjected those cells to RNA sequencing and gene expression analysis (Fig. 1B right).

**Figure 1:**
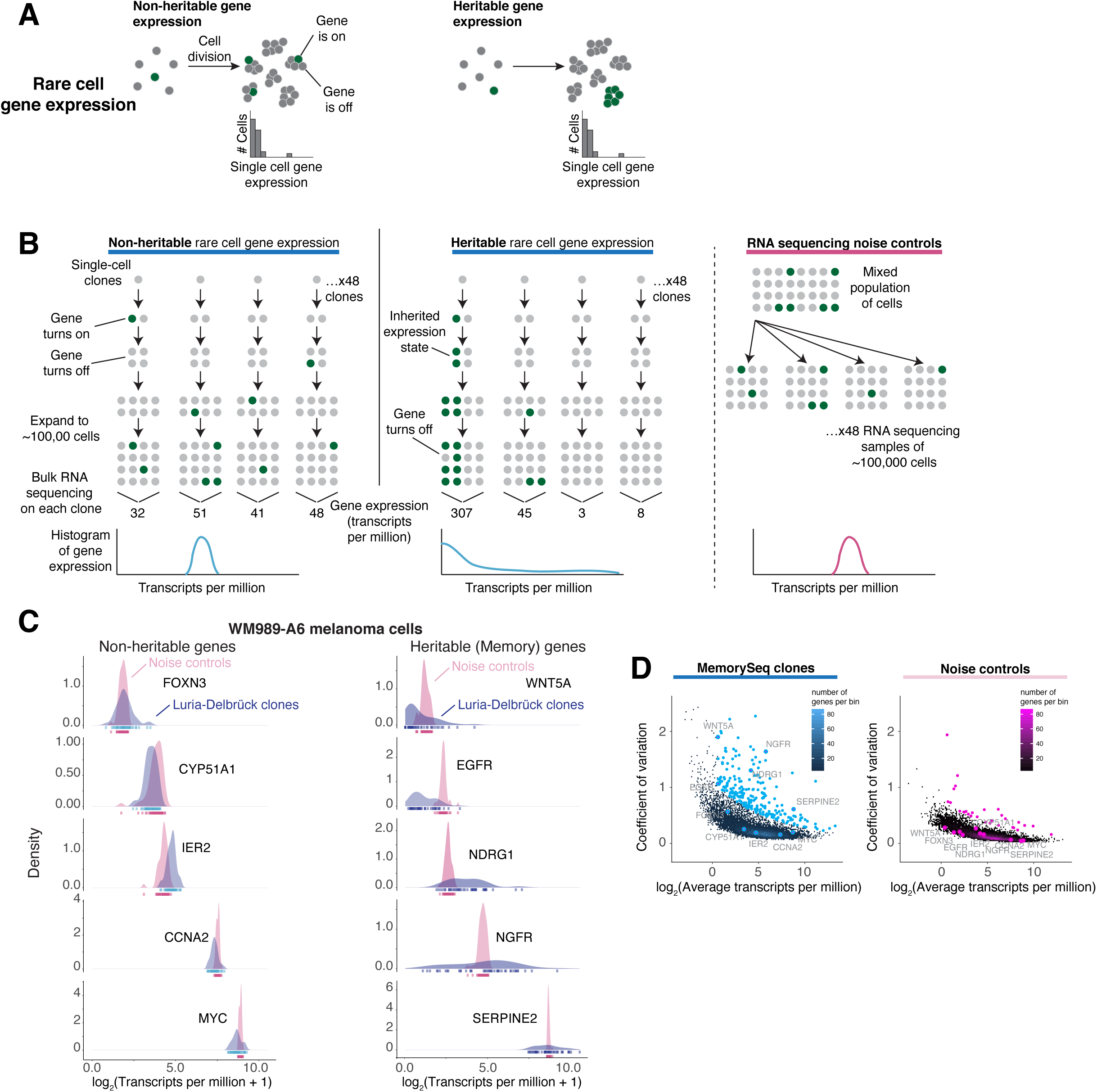
MemorySeq can identify genes with high transcriptional memory. **A.** Rare cell gene expression patterns, both with and without heritability. Histograms of single cell expression levels are unable to discriminate between these two alternatives. **B.** Schematic of MemorySeq experiment. We started with a single melanoma cell (WM989-A6), grew it to ∼100 cells, then seeded 48 wells from those cells and allowed the cells to proliferate to around ∼100,000 cells before subjecting the entire MemorySeq clone to RNA sequencing to determine expression levels. In the case of non-heritable expression, the levels of expression would not vary dramatically between MemorySeq clones, whereas in the heritable case, some clones would exhibit much higher levels of expression when a cell moved into the high expression level state early in the family tree of the clone. To determine how much variability in expression would arise for purely technical reasons, we also performed control experiments by plating around ∼100,000 cells directly into individual wells and performing RNA sequencing. **C.** Expression histograms across n=43 MemorySeq clones for genes identified as non-heritable (left) or heritable (right). **D.** Coefficient of variation *versus* mean expression levels for all 23,669 genes that we analyzed across all MemorySeq clones. Points labeled with blue dots (on MemorySeq clones plot) or pink dots (on Noise controls plot) passed the threshold for being identified as a heritable gene. These genes were identified by first fitting a Poisson regression model to the data, and then selecting genes with residuals in the top 2%. This approach identified 227 heritable genes from the MemorySeq clones, but only 30 genes passed this threshold in the Noise control condition. Particular genes from the panel in Fig. 1C are labeled on both plots.

We first applied MemorySeq to the melanoma cell line WM989-A6. We chose this cell line and culture system because we had already verified the presence of rare cells within the population marked by the expression of a particular subset of genes such as *EGFR, NGFR* and *AXL*. These rare cells were strongly associated with resistance to the targeted melanoma drug vemurafenib (Fallahi-Sichani et al., 2017; Shaffer et al., 2017), and the independent observation that sibling cells often expressed the same genes suggested that these genes displayed some degree of memory (Shaffer et al., 2017). Thus, in this system, we have already identified several genes that are both associated with a phenotype and appear to exhibit some degree of heritability. These genes naturally serve as positive controls for the MemorySeq methodology.

Upon performing MemorySeq in this cell line, we first checked the distribution of expression levels across MemorySeq clones for a number of previously identified resistance-associated genes and non-resistance associated genes (Fig. 1C, Heritable genes) (Shaffer et al., 2017). As hoped, we found that the resistance marker genes displayed far greater variability across clones than the technical noise controls. Conversely, housekeeping genes and other genes that do not exhibit much cell-to-cell variability showed variance across clones that was much more similar to that of the technical noise controls (Fig. 1C, Non-heritable genes). Intriguingly, *MYC*, a proto-oncogene for which we have seen high levels of cell-to-cell variability (Supp. Fig. 1), showed little increased variance across clones compared to controls, suggesting that its transcriptional memory was much lower than that observed in the marker genes (as verified by alternative means in Fig. 3C). We observed similar behavior for CCNA2, a cell cycle gene whose expression would similarly be expected to vary from cell to cell but not exhibit much heritability due to cell cycle desynchronization *(Chao et al., 2018)*; Fig. 3C.

**Figure 2:**
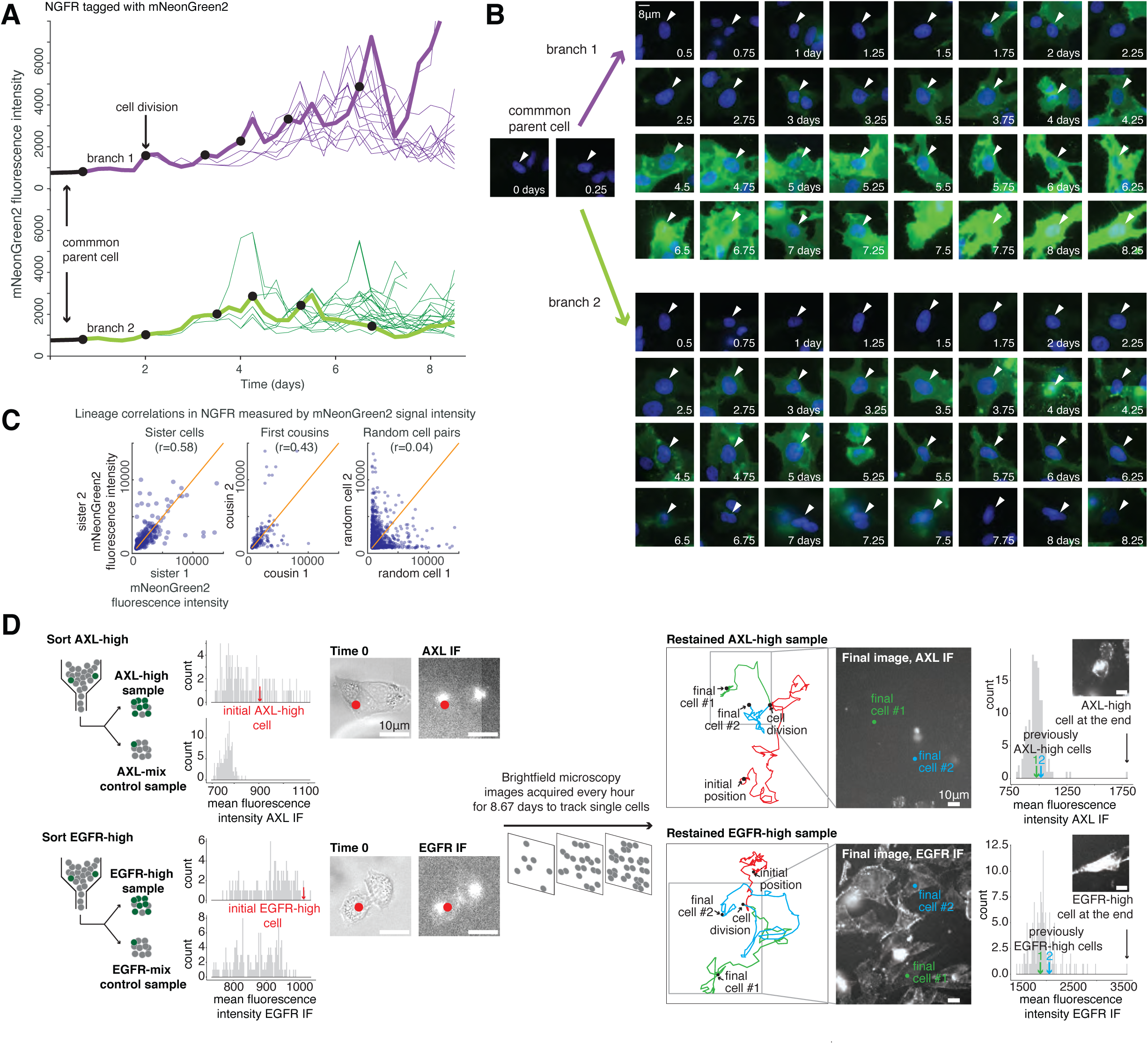
Time-lapse microscopy verifies rare, high expression states that persist for several cell divisions. We generated a cell line (WM989-A6-G3 C10-C2 clone E9) that expresses a large but incomplete (and thus nonfluorescent) portion of the mNeonGreen2 fluorescent protein with the remaining piece of mNeonGreen2 fused to NGFR at the endogenous locus. When the NGFR fusion protein expresses, the remaining portion of mNeonGreen2 binds to the NGFR fusion protein and becomes fluorescent. We then performed time-lapse microscopy imaging of the NGFR protein (nucleus labeled with H2b-iRFP670) at 6 hour intervals for 8.75 days. **A.** We tracked cells through several cell divisions, thus building cellular lineages, and quantified fluorescence intensity for each cell. The plot shows two branches from the same parent cell with fluorescence intensity of mNeonGreen2 over time. **B.** Series of fluorescent micrographs of the two cells highlighted in panel A. Scale bar is 8µm long. **C.** Correlations between sibling cells, first cousins, and random pairs of cells (n = 486, 292, 905, respectively). **D.** We stained cells with antibodies targeting AXL and EGFR, then sorted positive cells, plated them on a glass dish, and took images of their immunofluorescence signal. Subsequently, we acquired transmitted light images every hour for 8.67 days to facilitate tracking of cell lineages, and then we performed immunofluorescence again to measure EGFR and AXL levels at the end of the tracking period. From the timelapse images, we tracked selected lineages that initially contained cells with high levels of EGFR and AXL. The red dots on the left images correspond to the red arrow on the histogram for an example initial cell subjected to tracking. Upon division, we colored the tracks of the sibling cells green and blue respectively. The EGFR and AXL levels for these cells in their final state is indicated by the green and blue arrows on the histograms on the right. Scale bars are 10µm long.

**Figure 3:**
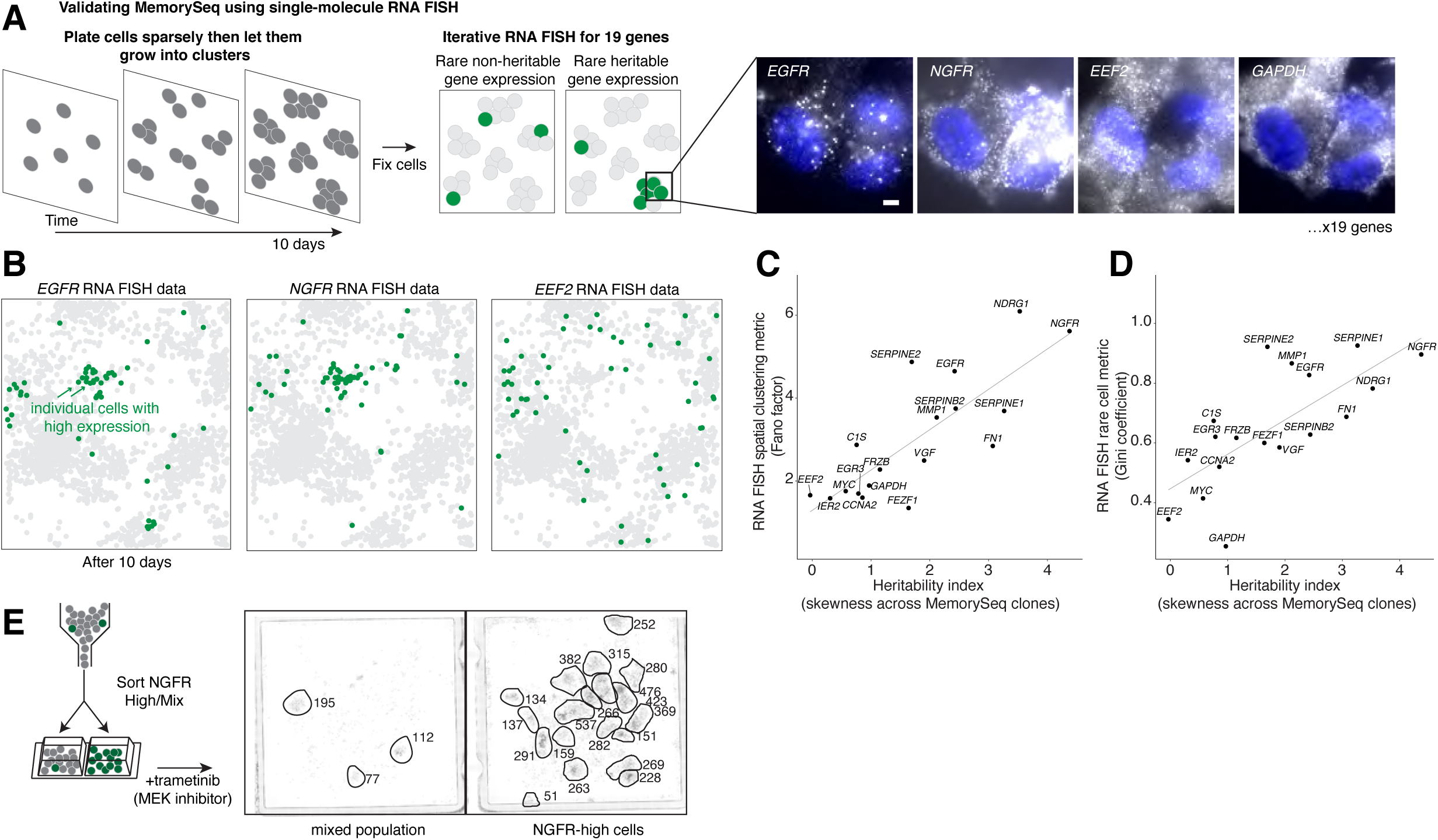
Single-molecule RNA FISH verifies the quantitative nature of MemorySeq for measuring heritability in single cells. **A.** Schematic of space-FISH experiment. We plated WM989-A6 melanoma cells sparsely on a dish and allowed them to grow for 10 days. We then fixed the cells and performed iterative RNA FISH to measure the expression of 19 genes. Closely related cells will remain in close proximity, thus heritable rare-cell expression would manifest as “patches” of *on* cells, whereas non-heritable rare-cell expression would display a more “salt and pepper” pattern of expression. Right: micrographs of RNA FISH for 4 genes, *EGFR* (heritable), *NGFR* (heritable), *EEF2* (housekeeping) and *GAPDH* (housekeeping). **B.** Each spot is a cell from an RNA FISH image scan of 12,192 cells (subset of 2,103 cells shown). Cells above a threshold (6 for *EGFR*, 36 for *NGFR* and 320 for *EEF2*) were considered to be in the high expression state and colored green. **C.** Quantitative comparison of heritability as measured by MemorySeq (x-axis: skewness across MemorySeq clones) and spatial RNA FISH analysis (y-axis). We used the Fano factor measured for spatial bins of 20 nearest cells as a spatial clustering metric; randomly placed high-expression-state cells would display a Poisson distribution and thus give a Fano factor of 1. Cell populations with a Fano factor greater than 1 would display some degree of spatial clustering. Of note, this plot and the plot in panel D contain 18 of the 19 genes that we quantified with RNA FISH because 1 gene (*CYR61*) did not pass the minimum mean transcripts per million cutoff for analysis in the MemorySeq data. **D.** We plotted MemorySeq heritability versus the Gini coefficient (from RNA FISH). The Gini coefficient measures expression inequality, and thus indicates the rareness of expression, with 0 being completely equal and 1 being completely unequal. **E.** Rare cells within clonal WM989-A6 populations marked by high levels of NGFR protein were sorted, cultured for 8-16 hours and then subjected to trametinib treatment at 10nM (MEK inhibitor) for 3 weeks. Image shows the number of resistant colonies (circled) along with number of cells within the resistant colony as indicated. Biological replicate in Supp. Fig. 14.

Given that RNA sequencing provides expression levels across the transcriptome, MemorySeq is able to measure heritability in the expression of all genes at once. Thus, we analyzed expression variance across clones for all genes. We found that for many genes across a range of average transcription levels, variance across clones was much higher than technical noise controls, suggesting that those genes exhibited high levels of transcriptional memory (Fig. 1D). We found that genes with higher expression levels typically had systematically lower variance across clones. By explicitly fitting this relationship, we could identify genes as high memory based upon their large residuals from the fit. We generated a panel of high memory genes with residuals in the 98th percentile or greater, and with a minimum expression level of 1.5 transcripts per million to eliminate spurious inclusion of lowly expressing genes, resulting in 227 genes identified as potentially having high heritability in WM989-A6.

Our experimental design predicted that a gene whose expression exhibits high variability across MemorySeq clones would occasionally initiate high levels of expression that would persist across multiple cell divisions but would not persist indefinitely. We sought to directly confirm these expression characteristics by using time-lapse microscopy to trace the expression state of individual cells for three genes. For one gene (*NGFR*), we were able to genetically tag the gene using a split-fluorescent protein approach (see Methods) to fuse mNeonGreen2(11) to *NGFR* (producing the NGFR-mNG2 protein). We used that cell line to track NGFR levels by fluorescence microscopy over a period of 8.75 days (Fig. 2; see Supp. Fig. 2 for cell line validation; single allele tagged, which may affect the levels of variability as compared to total protein). The vast majority of cells displayed essentially no fluorescent signal, but as predicted, occasional rare cells within the population displayed high levels of fluorescence. We then tracked 222 cell lineages through several cell divisions (examples of positive cells in Fig. 2A-B, Supp. Movie 1). We observed that cells would occasionally initiate high levels of expression of NGFR-mNG2 (compare top branch vs. bottom branch), and once initiated, that high level of expression could be maintained through multiple cell divisions, thus confirming the presence of memory. Further demonstrating the transience of this high expression state, we also observed cells transitioning from the high state to low levels of expression, with an average time in the high state of 40 hours; however, a few cells showed longer fluctuations in NGFR levels ranging from 4.25 to 5.25 days (Supp. Fig. 3; Supp. Fig. 4 for discussion of analysis).

Another prediction of long-lived but imperfect memory is that sibling cells should show greater correlation in expression levels than cousins. Confirming this prediction, we observed that the correlation of expression between recently divided sibling cells was higher (R=0.58) than between cousins (R=0.43), although both values were higher than that between unrelated cells in the population (R=0.04) (Fig. 2C). Demonstrating the phenotypic significance of these fluctuations in expression levels, we further found that cells expressing high levels of NGFR-mNG2 at the time of vemurafenib addition were much more likely to continue to proliferate (Supp. Fig. 5, Supp. Movie 2 and Supp. Movie 3).

We wanted to verify this same transient expression behavior for other genes, but it proved technically challenging to genetically tag genes such as *AXL* and *EGFR*. We thus used fluorescent antibodies to label AXL and EGFR, used FACS to isolate the high population, seeded that population on an imageable surface, and performed time-lapse microscopy on the transmitted light images in order to trace the lineage of these cells (Supplementary movie 4 for AXL, Supplementary movie 5 for EGFR). At the end of 8.67 days, we fixed the cells and performed immunofluorescence *in situ*, followed by imaging to measure AXL and EGFR levels. We were able to track a total of 53 cells (26 for *AXL*, 27 for *EGFR*) (originating from 4; 2 each for *AXL* and *EGFR*), of which we could confidently reidentify 29 after the second round of immunofluorescence. Of these, we observed that 15/15 and 6/14 of these cells starting from those that initially had high levels of AXL and EGFR (higher than all the negative cells; Fig. 2D), respectively, eventually turned off (<75th percentile) within the time window of 8.67 days (Fig. 2D and Supp. Fig. 6), demonstrating that the EGFR or AXL high state is indeed transient.

(We also verified this same transient expression behavior for other genes (*AXL, EGFR* mRNA, Supp. Fig. 7; NGFR protein, Supp. Fig. 8) by using FACS to isolate highly expressing cells and measuring the degree to which the expression levels in these cells reverted towards the distribution from the original population, finding a variety of timescales ranging from 5 to 9 days. Note that we observed an initial increase in expression for some of these genes upon sorting, which may be due to the stress associated with flow sorting or paracrine signaling in the concentrated subpopulation.)

While time-lapse microscopy provided direct evidence of the long-lived fluctuations predicted by MemorySeq, it is difficult to perform for a panel of genes owing to the challenges associated with editing genes, especially those whose expression is very low in most cells. Thus, we sought another method to confirm these heritable fluctuations for a larger panel of genes. First, we performed experiments in fixed cells grown on culture dishes to measure heritability in gene expression by using spatial proximity as a proxy for relatedness. We seeded cells sparsely in culture dishes and then allowed them to grow for approximately 10 days, at which point we fixed the cells and subjected them to iterative single-molecule RNA FISH to measure the expression of 19 genes with a range of MemorySeq values (see Supp. Fig. 9) in individual cells while preserving their spatial context (Fig. 3A; as in Ref. (Shaffer et al., 2017)). (Genes with high MemorySeq signals also displayed rare cell expression patterns as expected Supp. Fig. 10.) Our reasoning was that as cells divide, their spatial proximity would reflect their relatedness (Hormoz et al., 2016). In the case of a gene with non-heritable expression, one would expect to find no spatial correlation in which cells were deemed high expressing within the population. In contrast, for genes with heritable expression, one would expect to find the high expressing cells to appear in patches corresponding to related neighboring cells that share a common ancestor that transitioned to the high expression state. We found that genes identified by MemorySeq as being highly heritable (e.g. *EGFR, NGFR, NDRG1, SERPINE1*) tended to show patch-like expression patterns across large numbers of cells, confirming that their expression was indeed heritable. In contrast, genes that MemorySeq would predict to not be heritable exhibited a more salt-and-pepper (variable but not heritable) expression pattern, as expected (Fig. 3B; also Supp. Fig. 11).

We wondered to what extent MemorySeq could measure differences in heritability of the high expression state for different genes. We therefore compared the degree of heritability from MemorySeq (given by the amount of skewness in the distribution of expression across MemorySeq clones) to the degree of heritability from spatial RNA FISH analysis (given by the amount of patchiness in the population). We found a strong correspondence between these two metrics (adjusted R^2^ = 0.6193), suggesting that MemorySeq can stratify genes by the gradations in the degree of heritability that they display (Fig. 3C; see Supp. Fig. 12 for further analysis).

The timescales of particular genes turning between high (on) and low (off) expression states should in principle be quantitatively related to the measured variability across the MemorySeq clones, with high variability corresponding to slow switching and *vice versa*. We thus analyzed a stochastic model of cell proliferation and switching relating these two quantities (see Supp. Note 1). Under the assumption that the *on* state is relatively rare, this model yielded a direct relationship between the coefficient of variation measured by MemorySeq multiplied by the fraction of the time the cell is in the *on* state and the predicted memory (number of generations cells are on before turning off), with the only further parameter being the total number of divisions in the MemorySeq experiment. This equation predicted that over a relatively large range of reasonable parameters, (CV ∼ 0.5-2, fraction of time on ∼ 0.01), the predicted memory was mostly in the range of 5-10 divisions, matching our experimental observations.

Motivated by our previous work in this cell line, we also looked for correspondences between the degree of heritability and the rarity of gene expression as measured by the Gini coefficient, a metric for inequality (Jiang et al., 2016; Shaffer et al., 2017; Torre et al., 2018). We observed that indeed the two metrics were correlated (adjusted R^2^ = 0.4898) (Fig. 3D), suggesting that heritable genes identified by MemorySeq are more likely to express only in rare cells. This correspondence may be due to the design of the MemorySeq experiment, in which skewness can reach potentially higher levels for rarer expressing genes than for less rarely expressing genes.

Having validated that MemorySeq was accurately identifying genes displaying transcriptional memory, we then asked what these genes were and what their expression in rare cells signified. The underlying hypothesis was that these slow fluctuations are more likely to be associated with distinct cellular behaviors in those cell subpopulations than fast fluctuations. Our reasoning was that a distinct cellular behavior would likely require a persistently different gene expression pattern, involving deviations in the expression of several genes simultaneously, as opposed to a transient (and, as we hypothesized, probably inconsequential) fluctuation. In this melanoma cell line, we have previously shown that rare cells have high levels of expression of certain genes associated with therapy resistance (including *EGFR, NGFR*, and *AXL*), and that these rare cells are much more likely to survive the initial application of drug to develop into resistant clusters. Thus, we first wondered whether the set of genes identified by MemorySeq that mark rare cells overlapped with the set of genes associated with resistance. We found that most MemorySeq heritable genes (162 out of 227) were also markers of resistance (as determined by Shaffer *et al.*, Supp. Fig. 13). These results suggest that the genes identified by MemorySeq are expressed in cell populations that are phenotypically distinct from the bulk of the population.

To verify the phenotypic differences of these cells, we used FACS to isolate cells by either high levels of NGFR or EGFR expression, and then we subjected them to a targeted inhibitor of MEK, trametinib, used to treat melanoma. On unsorted populations, upon treatment with this drug, a small percentage of cells will continue to grow and form colonies, mimicking the acquisition of drug resistance. In congruence with previous results, the EGFR/NGFR-high subpopulations resulted in far more resistant colonies after application of the drug, showing that this subpopulation is highly enriched for pre-resistant cells (Fig. 3E; Supp. Fig. 14). This result demonstrated that MemorySeq revealed the same subset of cells that we had previously determined to be highly enriched for drug resistant cells.

Our results thus far highlight MemorySeq’s ability to prospectively reveal functionally distinct subpopulations within clonal populations of apparently homogeneous cells. In the case of the WM989-A6 melanoma cell line, we had already established the existence of such a subpopulation, but for most cell lines, there is little to no information about single-cell fluctuations that exhibit memory and thus may also be associated with distinct phenotypes. We thus set about testing MemorySeq in another cell line, MDA-MB-231-D4, which is a clonal derivative of a triple negative breast cancer cell line (does not express *HER2*, estrogen or progesterone). Paclitaxel is a drug used to treat such breast cancers, but while it is able to kill most MDA-MB-231-D4 cells, some cells in the population are still able to survive the drug, thus leading to drug resistance (Fig. 4A). However, prospective markers to isolate the subpopulation of cells that are resistant to drug have remained elusive (Gao et al., 2017), and we hypothesized that MemorySeq might be able to reveal genes whose expression was associated with this single cell phenotype.

**Figure 4:**
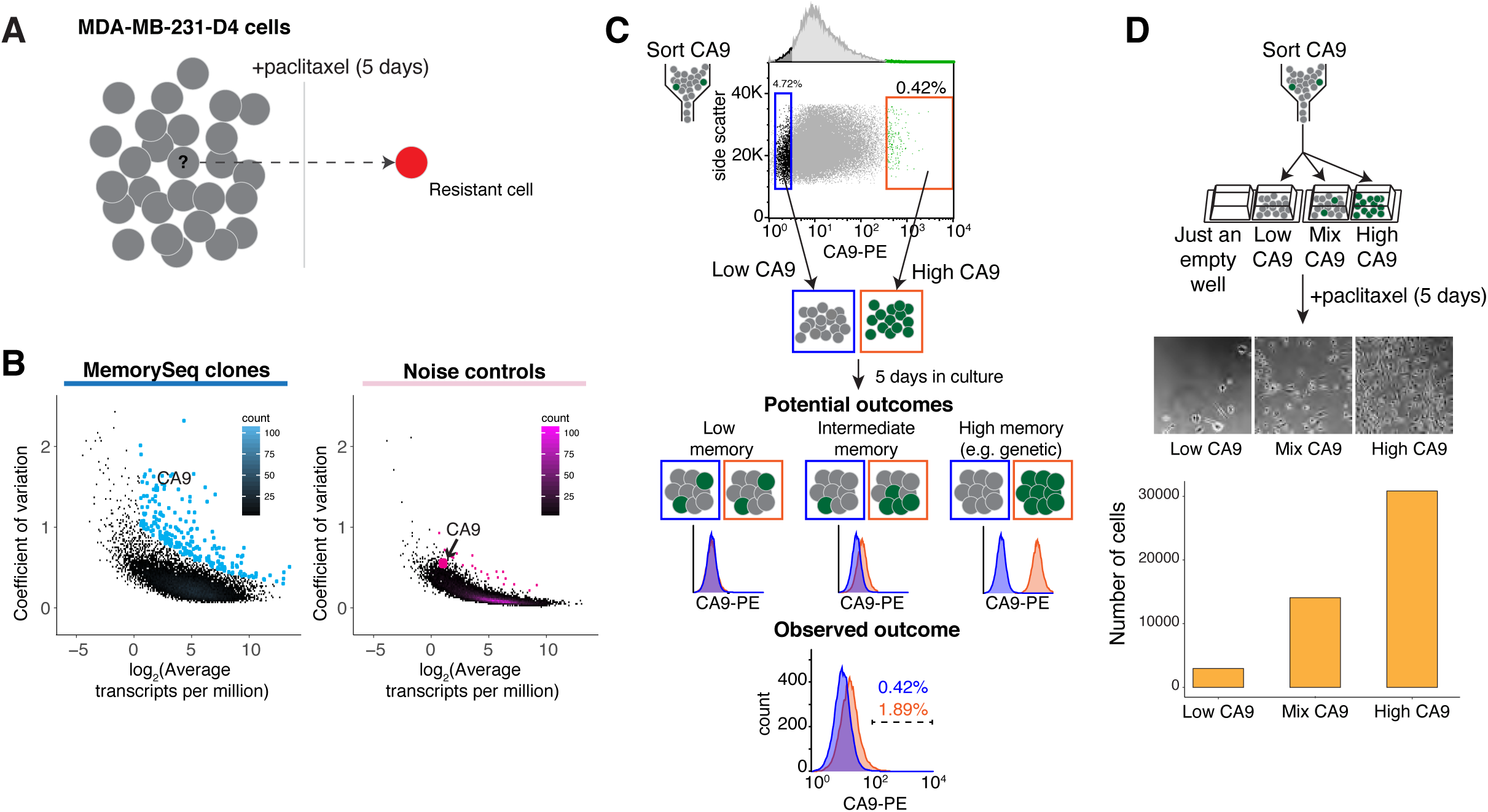
MemorySeq reveals a rare subpopulation of MDA-MD-231-D4 cells associated with drug resistance. **A.** Most MDA-MD-231-D4 cells die upon treatment with paclitaxel for 5 days, but a small subpopulation of cells (cell marked with “?”) survive and become resistant (red cell). **B.** We performed MemorySeq analysis on MDA-MD-231-D4 cells (n=39 clones, left; n=46 control clones, right). The blue colored dots correspond to genes that we statistically identified as being highly heritable by fitting a Poisson regression model and selecting genes with residuals in the top 2%, as was done with WM989. **C**. We stained cells with antibody targeting the CA9 surface marker and then sorted out the top 0.5% of cells and the lowest 5%. After culturing for 5 days, we re-stained the cells and measured CA9 levels by flow-cytometry. Potential outcomes for the levels of CA9 staining depending on the degree of gene expression memory. Observed outcome below for CA9 staining 5 days after initial sorting. **D.** We stained cells with antibody targeting the CA9 surface marker and then sorted out the top 2-4% of cells, the lowest 2-4% of cells, and the total “mix” population into chamber wells, after which we applied paclitaxel 1 day after sorting for 5 days. Transmitted light micrographs show the number of cells remaining after drug treatment for the different populations, and the quantification of the number of cells was performed using cell counting based on nuclear identification by imaging the DAPI nuclear stain and identifying computational techniques. All scale bars are 50µm long.

To test this hypothesis, we performed MemorySeq on the MDA-MB-231-D4 cell line by growing each of 48 subclones to around 100,000 cells, after which we performed RNA sequencing and quantification as described for WM989-A6. As with the melanoma cell lines, MemorySeq revealed a large number of genes in the MDA-MB-231-D4 cell line, with putative heritable expression patterns (230 genes, Fig. 4B). The range of variability in expression levels for all genes across the MemorySeq clones (most genes having low variability, but a few showing high variability) was comparable to that identified by MemorySeq for the WM989-A6 cell line. Interestingly, however, the overlap between the gene sets was relatively small (Supp. Fig. 15), suggesting that different cell lines may have distinct sets of “memory genes” and that there is no universal “memory gene” expression program (with a potential exception noted below). As with WM989, we confirmed both the rarity and heritability of the expression pattern by using RNA FISH on cells initially seeded sparsely and then allowed to grow in place for 10 days, as done for the melanoma lines (Supp. Fig. 16, compare to Fig. 3C).

Given the existence of these rare, slowly fluctuating subpopulations in the MDA-MB-231-D4 cells, we next asked whether these newly identified subpopulations were associated with phenotypic differences such as differential sensitivity to paclitaxel. Amongst the genes identified by MemorySeq was CA9, a surface marker known to be negatively associated with breast cancer chemosensitivity (Aomatsu et al., 2014; Span et al., 2003), but for which there was no reason to suspect that its expression at the single cell level would be indicative of which cells specifically survived upon drug treatment. We thus immunolabeled the MDA-MB-231-D4 cells using antibodies targeting CA9 and then used FACS to isolate a rare population of CA9 positive cells (2-4% percent, along with CA9 low and mixed subpopulations; validation of sorting in Supp. Fig. 17), after which we added paclitaxel to both and grew the cells for 5 days. We found that, when treated with 1nM paclitaxel, the CA9 positive cells were more likely to be resistant than either the CA9 low or mixed subpopulations. (Furthermore, the sorted subpopulations reverted to the population average, demonstrating that the CA9-high state is ultimately transient; Supp. Fig. 18.) These results, in a cancer cell line of a completely different type involving a drug with a completely different mechanism of action, demonstrate that MemorySeq is able to identify *de novo* heritable, rare cell expression states, and that these rare cells are phenotypically distinct from the others in the population.

Behavioral differences such as drug resistance are typically associated with the differential expression of many genes at once. We thus further hypothesized that the long timescale of these single cell fluctuations could allow for significant co-fluctuation; that is, if a cell expresses a high level of one high memory gene for a sufficiently long time period, it could also have a higher probability of expressing another slowly fluctuating gene simultaneously. Indeed, should such a phenomenon be prevalent, it would allow us to organize these high memory genes into characteristic modules of genes that co-fluctuate in single cells.

To isolate such modules, we calculated the correlation coefficient between the expression of all pairs of heritable genes across the MemorySeq clones derived from WM989-A6. We reasoned that if a particular clone had a high abundance of a particular transcript, then the abundance of transcripts of co-fluctuating genes would also be high in that particular clone. We saw large blocks of genes whose expression appeared to correlate strongly with each other, suggesting that they co-fluctuate at the single cell level (Fig. 5B). To validate that the programs so identified by MemorySeq corresponded to single cell correlations, we compared the correlations between MemorySeq and RNA FISH in single cells on a panel of genes across two separate clusters. We found a strong general correspondence between these two assays, suggesting that MemorySeq is able to identify groups of genes *de novo* that co-fluctuate in rare-cell expression programs (Fig. 5C, Supp. Fig. 19 and Supp. Fig. 20). We observed similar clustering in another melanoma cell line (WM983B-E9; see Supp. Fig. 21) and MDA-MB-231-D4, although the specific genes were typically different (Supp. Fig. 15). We also performed MemorySeq on the lung cancer cell line PC-9, which showed a total of 240 heritable genes, including 8 genes that were heritable in all of the other cell lines on which we performed MemorySeq (WM989, WM983B, and MDA-MB-231, Supp. Fig. 22).

**Figure 5:**
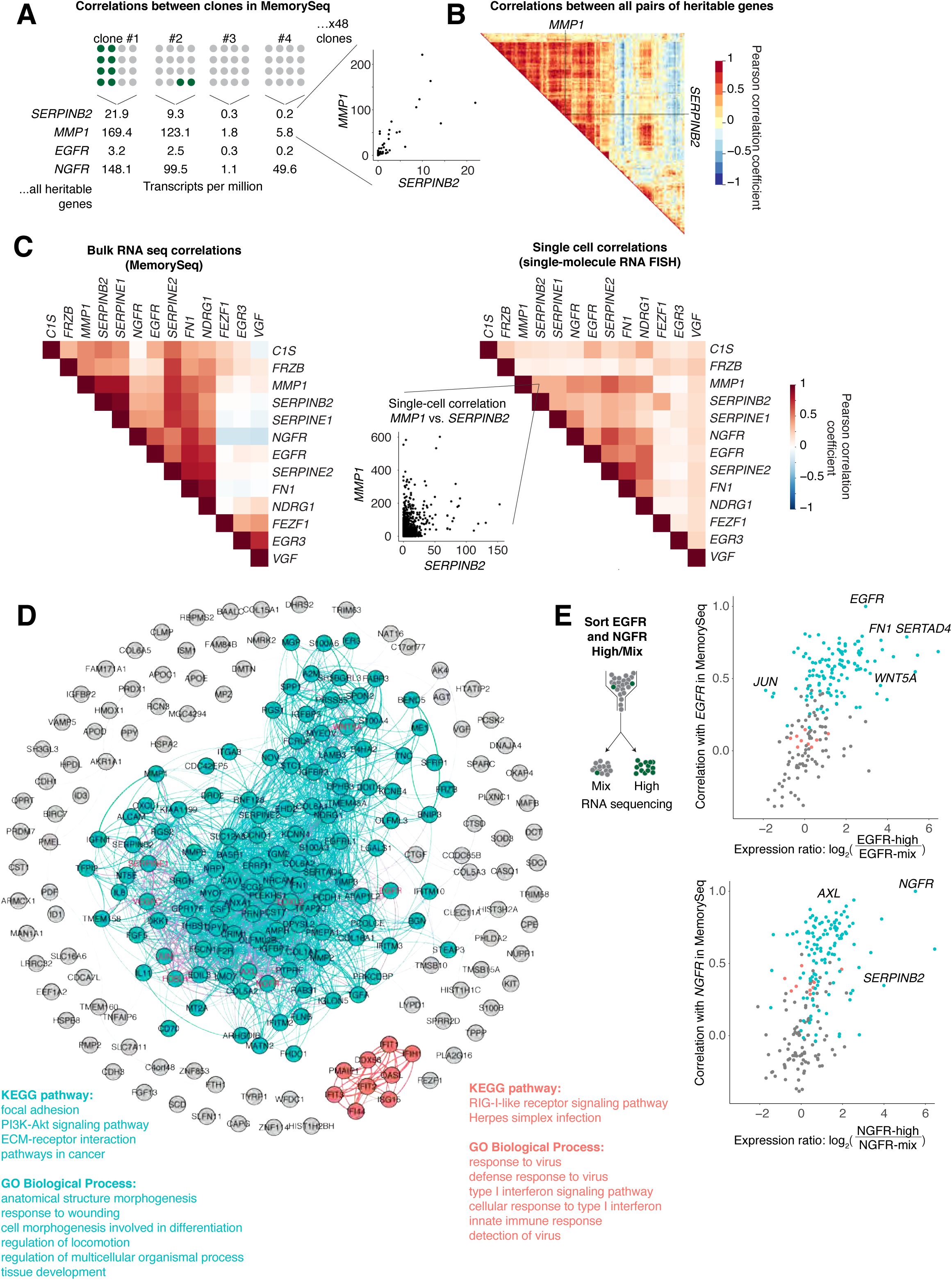
MemorySeq enables the identification of coordinated rare-cell expression programs. **A.** We measured correlations between genes across MemorySeq clones derived from WM989-A6 melanoma cells. Shown is an example correlation between *MMP1* and *SERPINB2* across 43 MemorySeq clones. **B.** Correlations between all pairs of genes exhibiting heritability as determined by the threshold described in Fig. 1. Cook’s distance analysis to test for outliers is given in Supp. Fig. 25. **C.** Comparison of coherence between MemorySeq bulk RNA-seq analysis and single cell correlations as measured by single-molecule RNA FISH. We performed RNA FISH on 20 genes in WM989-A6 cells, keeping for further analysis genes whose RNA FISH Gini coefficient was greater than 0.6 (13 genes remaining). The correlation between bulk MemorySeq RNA-seq levels is on the left, RNA FISH on the right. Callout shows raw RNA FISH counts for 12,192 cells between *MMP1* and *SERPINB2* in single cells. **D.** Community detection within the network defined by the correlation matrix of co-expression patterns among the heritable genes. Gray circles indicate genes that did not comprise a network community. Green and Red indicate the two communities detected; KEGG pathway and GO Biological Process analysis results shown for both communities. **E.** Comparison of rare cell expression programs identified by MemorySeq and those identified by sorting EGFR (top) or NGFR (bottom) high cells (using fluorescent antibody labeling followed by RNA-seq on the high versus mix populations).

The appearance of distinct clusters of slowly co-fluctuating genes led us to use community detection algorithms for network data to demarcate these groups of genes for further analysis. We used a weighted version of *k*-clique community detection (Derényi et al., 2005; Jonsson et al., 2006; Palla et al., 2005) to identify such groups of genes (Fig. 5D) (see Methods for more information). We chose *k*-clique community detection because it allows for nodes to be in multiple communities at once, echoing the ability of a protein to simultaneously play multiple roles within the cell. In WM989-A6 cells, one large community of genes overlapped very strongly with the vemurafenib pre-resistance marker gene set that we identified earlier (Shaffer et al., 2017). We verified this correspondence by comparing the transcriptomes of sorted subpopulations of EGFR-high cells to the clusters of genes identified by MemorySeq, showing that the expression levels of genes specifically in this cluster correlated well with that of genes that correlated with EGFR expression (Fig. 5E). (All networks shown in figures were visualized with Gephi (Bastian et al.).)

We also found other communities within the set of heritable genes in WM989-A6 cells, suggesting the existence of multiple independent heritable gene expression programs. One prominent one included *DDX58* (RIG-I), *IFIT1, PMAIP1*, and *OASL*, which may be related to type 1 interferon signaling (Loo and Gale, 2011), and was notable because it also appeared to some extent in MDA-MB-231-D4 and WM983B-E9 (Supp. Fig. 23). We verified that this cluster expressed in a distinct rare-cell subpopulation from that containing *EGFR* and *AXL* by performing RNA FISH for *PMAIP1, DDX58, AXL*, and *EGFR* simultaneously. This analysis showed that *PMAIP1* and *DDX58* expression exhibited very strongly correlated rare-cell expression, but neither correlated to much extent with either *AXL* or *EGFR* expression (Supp. Fig. 24). This cluster appeared relatively distinct from the primary community associated with drug resistance and *EGFR* expression, and indeed, did not show any association with the EGFR-high transcriptome (Fig. 5E). Another community in WM989-A6 cells was somewhat less coherent, and included genes such as *VGF. VGF* expression also showed strong heritability (Fig. 3C) but its expression levels appeared not to correlate with the other pre-resistance genes at the single cell level (Fig. 5E), and was not associated with the drug resistance phenotype (Shaffer et al., 2017).

The fact that multiple genes appeared to coordinate their expression across multiple chromosomes suggested that the mechanism for maintaining memory occurs in *trans*, i.e., through the regulatory milieu rather than just a short but intense pulse of transcription that propagates to children through partitioning upon cell division. Labeling sites of nascent transcription for EGFR (Levesque and Raj, 2013) revealed that transcriptional activity also occurred in patches (as opposed to just transcript abundance), further suggesting that memory of the high expression state was due to sustained transcriptional activity as opposed to a single sporadic, large, transient burst of transcription in a precursor cell (Supp. Fig. 25 and Supp. Fig. 7).

In sum, we believe MemorySeq is a simple but powerful method for identifying rare, heritable expression patterns in cells. We have shown here that such rare, heritable expression programs may be related to non-genetic mechanisms of therapy resistance in cancer (Pisco and Huang, 2015; Roesch et al., 2010; Shaffer et al., 2017; Sharma et al., 2010; Spencer et al., 2009).

However, they may also be important in other contexts in which rare cells behave differently than the rest of the population, both in cancer (such as metastasis), but also in otherwise healthy tissues or in cellular reprogramming contexts like the induction of iPS cells (Hanna et al., 2009).

It is also interesting that MemorySeq is quantitative in the sense that it is not just able to report that a gene’s expression is heritable, but is also able to provide a relative sense of how heritable that expression is, meaning how many cell divisions does it require before cells begin to forget the rare-cell expression state. By revealing such intermediate timescales of fluctuation, the results of MemorySeq suggest that the classic use of the concept of somatic epigenetics (non-genetic heritability) may require reevaluation as a quantitative, rather than qualitative, property of some cellular states.

## Materials and Methods

### Cell lines and culture

We used 3 cell lines in our study: WM989-A6, which is a subclone of the melanoma line WM989 (Wistar Institute, a kind gift of Meenhard Herlyn); WM983B-E9, a subclone of WM983B (Wistar Institute, a kind gift of Meenhard Herlyn); and MDA-MB-231-D4, a subclone of MDA-MB-231 (ATCC HTB-26). We verified all cell lines by DNA fingerprinting: WM989-A6 and WM983B-E9 were performed at the Wistar Institute by DNA STR Microsatellite testing and the MDA-MB-231 was performed by ATCC human STR profiling cell line authentication services). We cultured all melanoma cell lines in the recommended medium, which is TU2% for the melanoma lines (containing 80% MCDB, 10% Leibovitz’s L-15, 2% FBS, and 2.4mM CaCl_2_) and DMEM with 10% FBS for MDA-MB-231.

To create the NGFR-mNG2(11) WM989-A6-G3 cell line, we used the split mNeonGreen2(mNG2) system described in Ref. (Feng et al., 2017). In brief, we first transduced WM989 A6-G3 with 10/11ths of mNG2, which is non-fluorescent without the remaining 1/11th of the protein. We then electroporated cells with Cas9 RNP containing a guide RNA specific to the C-terminus of *NGFR* and a single-stranded DNA template encoding the remaining 1/11th of the protein flanked by sequences homologous to the targeted locus (sequences available on Dropbox in https://www.dropbox.com/s/je51xsok1ej2xtc/MemSeq_NGFR_NG2_Sequences.xlsx?dl=0). We then isolated fluorescent cells by FACS and generated clonal cell lines by serial dilution. To verify in-frame integration of the mNG2(11) construct, we PCR amplified the C-terminus *NGFR* locus from cell lysates and cloned the amplicon into a recipient plasmid. Half of the resulting plasmids contained the in-frame mNG2(11) sequence and the remaining half contained the unedited *NGFR* sequence without substitutions, insertions or deletions. Sanger sequencing traces are located are located here: https://www.dropbox.com/sh/65rvfs3pka5zii4/AAAwBZOynkB4rU9NJvCo_RJha?dl=0. We further confirmed that mNG2 fluorescence correlates with NGFR mRNA abundance at the single-cell level by single-molecule RNA FISH and validated that the NGFR-mNG2(11) WM989-A6-G3 cell line recapitulates phenomenology described in Shaffer *et al.* (2017), showing increased resistance to vemurafenib in the mNG2-high cells (Supp. Fig. 2; experiments described in RNA FISH, FACS, and drug treatment methods below). We also demonstrated that the localization of the protein was identical to that of the endogenous protein by comparing the NGFR-mNG2 fluorescence to that of the signal produced by immunofluorescent labeling of NGFR (Supp. Fig. 2). To facilitate cell tracking in the time-lapse images, we transduced cells lines in which we tagged NGFR with mNG2(11) with lentivirus encoding an H2B-iRFP670 which localizes to the nucleus, thus enabling us to track cell nuclei. Following transduction, we derived clonal cell lines by serial dilution before imaging. All plasmid sequences are available here: https://www.dropbox.com/sh/wpgyfup6gwiyaeb/AAC0gRgg0dmBJoUlypvSDWnIa?dl=0.

To validate that cells with high levels of NGFR-mNG2 (as measured by fluorescence) were more resistant to vemurafenib, we trypsinized and pelleted the NGFR-mNG2(11) WM989-A6-G3 cell line, washed cells once with PBS containing 2mM EDTA, then resuspended in PBS containing 2mM EDTA and 100ng/mL propidium iodide or 200ng/mL 7-AAD and proceeded with sorting. Using a MoFlo Astrios (Beckman Coulter) or FACSJazz (BD Biosciences), we isolated the top 0.5-1% of mNG2 fluorescent cells and equal numbers of the bulk population gated only for live cells. We then treated these samples with vemurafenib as described in the drug treatment methods below. All flow cytometry data is available on Dropbox here: https://www.dropbox.com/sh/9bq1eg0k5o0q452/AACunY5g1xtp5lxSIox1OqOka?dl=0.

### MemorySeq

Our experiment roughly followed the design of Luria and Delbrück’s original fluctuation analysis, but with RNA sequencing as the terminal readout instead of the number of resistant colonies. From the parental cell line (WM989-A6, WM983B-E9, MDA-MB-231-D4), we isolated a single cell, let it proliferate until reaching roughly 100-200 cells, then plated these cells into a 96 well plate at a dilution in which we expected roughly 0.5 cells per well. From these wells, we isolated ∼100 clones for further expansion, excluding wells that were seeded with more than 1 cell. Of the 100 starting clones, we aimed for 48 clones from each cell line for downstream analysis. We grew the clones until they reached a minimum of around 100,000 cells, with some reaching as high as roughly 200,000 cells. At that point, we used miRNAeasy RNA extraction kit (Qiagen 217004) to isolate RNA from each clone, followed by library preparation using the NEBNext Poly(A) Magnetic Isolation Module and NEBNext Ultra RNA sequencing library prep kit for Illumina (NEB). At the time of RNA isolation for the clones, we also isolated 48 separate samples of 100,000 cells from the parental line and prepared these samples for RNA sequencing as controls. For each cell line, we sequenced a total of 96 samples, including 48 clones and 48 controls from a mixture of the parental cell line. We sequenced to a depth of at least 500,000 reads per RNA sequencing library (with a typical depth of around 4 million reads) on a NextSeq500 (Illumina). While we targeted 48 clones and 48 controls for each cell line, we had a few samples with poor RNA quality and occasionally lost samples when preparing the libraries. Therefore, after culturing, extracting RNA, and preparing libraries, we ended up with 39-46 clones total for each cell line for our analysis (43 clones for WM989, 46 clones for WM983B, and 39 clones for MDA-MB-231). We aligned the reads using STAR and enumerated read counts per gene using HTseq (code available at https://bitbucket.org/arjunrajlaboratory/rajlabseqtools).

For computational analysis of the Luria-Delbruck RNA sequencing data, we calculated the transcripts per million of every gene in each sample. We then calculated metrics of the variation across the different 48 clonal samples, including the coefficient of variation, skewness, and kurtosis. We also compared these metrics in the clonal samples to those observed in the mixed controls. We found that the relationship between the coefficient of variation and the transcripts per million for every gene could be fit by a Poisson regression model. We fit this model for each cell line and then defined the panel of heritable genes as those with residuals greater than the 98th percentile. We also set a minimum level of expression for heritable genes as 1.5 transcripts per million for WM989 and MDA-MB-231 and 1.5 transcripts per million for WM983B. This approach yielded 227 heritable genes in WM989, 230 heritable genes in MDA-MB-231, and 230 heritable genes in WM983B. We generated correlation matrices from the pairwise Pearson correlation coefficients for heritable genes across all clones. We calculated the Cook’s distance for the pairwise correlations to determine sensitivity to outliers (Supp. Fig. 26). For a few pairs, the correlation coefficient was deemed sensitive to the presence of outliers. This sensitivity is to be expected because of the experimental design; as per Luria and Delbrück’s fluctuation analysis, there will be rare outlier cultures. Given that we did not observe such outlier cultures in our technical controls, we believe that these outlier cultures reflect true biological variability. Computational analysis of RNA-sequencing data is available on the Dropbox here https://www.dropbox.com/sh/7z0n6ixshghdvlo/AADb7cY9ZGFMF1CRbLJpnYQSa?dl=0 and https://www.dropbox.com/sh/vykm0gu0afy39dm/AAAjPMibwX5LfaPWHnpD5k67a?dl=0. Lists of heritable genes are in Supp. Table 1.

### Fluorescence Activated Cell Sorting (FACS)

We stained WM989 A6-G3 melanoma cells for fluorescence assisted cell sorting using antibodies for EGFR and NGFR. We note that while we stained for both proteins, we did not isolate enough EGFR-high cells for testing trametinib resistance in all three replicates. First, for EGFR staining, we trypsinized 40-50 million cells, washed once with 0.1% BSA-PBS, and incubated for 1 hour at 4°C with 1:200 mouse anti-EGFR antibody, clone 225 (Millipore, MABF120) in 0.1% BSA-PBS. Next, we washed with 0.1% BSA-PBS and then incubated for 30 minutes at 4°C with 1:500 donkey anti-mouse IgG labeled with Alexa Fluor 488 (Jackson Laboratories, 715-545-150). We washed the samples again with 0.1% BSA-PBS and then incubated for 10 minutes at 4°C with 1:250 anti-NGFR antibody conjugated directly to PE-Cy7 (Biolegend, 400126, clone ME20.4) in 0.5% BSA-PBS with 2mM EDTA. Finally, we washed the samples with 0.5% BSA-PBS containing 2mM EDTA, then resuspended in 1% BSA-PBS containing 100ng/mL propidium iodide or 200ng/mL 7-AAD and proceeded with sorting using a MoFlo Astrios (Beckman Coulter) or FACSJazz (BD Biosciences). To aid with gating, we incubated control samples without the anti-EGFR primary antibody and with a PE/Cy7 mouse IgG1 isotype control (Biolegend, 400126). After gating for live cells, we collected either the top 0.02-0.2% EGFR-high cells or the top 0.5% NGFR-high cells. We also collected equal numbers of the bulk population by using the same gating for live cells, but without gating on either the EGFR or NGFR stains.

To monitor the dynamics of *AXL* expression, we stained WM989-A6-G3 melanoma cells for fluorescence assisted cell sorting by trypsinizing 40-50 million cells, washing once with 1% BSA-PBS, and incubating for 30 minutes at 4°C with 1:50 goat anti-AXL antibody (Novus AF154, lot DMG0516031) in 1% BSA-PBS. Next, we washed the cells twice with 1% BSA-PBS and then incubated for 30 minutes at 4°Cwith 1:85 bovine anti-goat IgG labeled with Alexa Fluor 647 (Jackson Laboratories, 805-605-180). Finally, we washed the samples with 1% BSA-PBS, then resuspended in 1% BSA-PBS containing 100ng/mL propidium iodide or 50ng/mL DAPI and proceeded with sorting. After gating for live cells, we collected the top 0.5-1% AXL-high cells and equal numbers of the lowest 80%-95% AXL-low cells, then plated cells onto glass-bottom chamber plates. After 1, 3, 6 or 9 days in culture, we fixed the sorted cells for RNA FISH as described below. To account for cell growth and changes in cell density, we plated fewer cells for later time-points. We performed a similar set of experiments for *EGFR* and *NGFR* using the staining procedure described above. For *NGFR*, we re-stained the sorted population after 7 days in culture (following the same procedure described above) and assessed NGFR intensity by flow cytometry using the same instrument as the initial sort.

To monitor the dynamics of *CA9* expression in MDA-MB-231 cells, we trypsinized cells, washed once with 0.1% BSA-PBS then stained with anti-CA9 antibody conjugated to phycoerythrin (PE) (Miltenyi Biotech clone REA658, 130-110-057) at a dilution of 1:11 in 0.1% BSA-PBS for 30 minutes at 4°C. After staining we washed the cells twice with 0.1% and resuspended in 0.1% BSA-PBS containing 200ng/mL 7-AAD and proceeded with sorting. After gating for live cells, we sorted the top 0.5%-2% CA9-high and the bottom 5-10% CA9-low cells. For 2 of 3 replicates, we stained the sorted cells with 2.5-5µM CellTraceViolet (Invitrogen C34557) in PBS at 37°C for 20 minutes, followed by 2 washes with media. We then culture the cells for 5 days before re-staining the cells and measuring CA9-PE and CellTraceViolet intensities by flow cytometry.

For testing the response of CA9-high cells to paclitaxel, we stained MDA-MB-231 cells as described above then sorted the top 2-3% of CA9-high, the bottom 2-3% CA9-low, and a mixed population using only the live cell gates (CA9-mix). After allowing the sorted cells to adhere overnight, we began treatment with 1nM paclitaxel as described below. We performed single-molecule RNA FISH for CA9 confirming that sorting with CA9 antibody enriched for CA9-high expressing cells (Supp. Fig. 17).

### Live-cell immunofluorescence

We stained and sorted AXL-high and EGFR-high cells as described above, then proceeded with live-cell imaging on a Nikon Ti-E inverted microscope enclosed in a temperature controlled and humidified chamber at 5% CO_2_. We acquired an initial set of images measuring Alexa Fluor 488 and Alexa Fluor 647 fluorescence to verify that the cells indeed had high protein levels, then proceeded to scan the slide (20x magnification) in brightfield every hour for 8 days and 16 hours. We then incubated the live cells with 1:80 goat anti-AXL antibody or 1:200 mouse anti-EGFR antibody in TU2% for 1 hour at 4°C followed by two washes with TU2%, secondary incubation with 1:250 bovine anti-goat Alexa647 or 1:250 donkey anti-mouse Alexa488, and two final washes with TU2%. After this re-staining, we imaged these live cells for Alexa Fluor 488 and Alexa Fluor 647 fluorescence at 20x magnification and quantified fluorescence intensity using rajlabimagetools available: https://bitbucket.org/arjunrajlaboratory/rajlabimagetools/wiki/Home and custom MATLAB scripts available at https://www.dropbox.com/sh/zrc0g9vtxewcctl/AABVYCaXHYyvWFPOx4ipXLJGa?dl=0

### Drug treatment experiments

We made stock solutions of 4mM trametinib (GSK1120212, Selleckchem, S2673), 4mM vemurafenib (PLX4032, Selleckchem, S1267), and 4mM paclitaxel (LifeTechnologies, P3456). For drug treatment experiments, we diluted the stock solutions in culture medium to a final concentration of 10nM for trametinib, 1µM for vemurafenib, and 1nM for paclitaxel. For trametinib treatment experiments, we sorted WM989-A6 by NGFR levels as described in the FACS section of Methods and then cultured them for 2-3 weeks. For vemurafenib experiments, we cultured the FACS sorted NGFR-mNG2 WM989-A6-G3 in vemurafenib for 21 days. For paclitaxel experiments, we cultured CA9 FACS sorted MDA-MB-231 cells in paclitaxel for 5 days. At the end of all drug treatment regimens, we fixed each sample in 4% formaldehyde for 10 minutes, permeabilized the sample in 70% ethanol, and then performed cell quantification.

### Cell quantification

We quantified cell numbers for drug treatment experiments by fixing the cells, staining with DAPI, then imaging across the majority of the well via image scanning at 20x magnification. After scanning, we computationally stitched the images together, after which we used custom software written in MATLAB to identify nuclei, which is publicly available here: https://bitbucket.org/arjunrajlaboratory/colonycounting_v2/wiki/Home.

### Time-lapse imaging

We acquired time-lapse images of the NGFR-mNG2 WM989-A6-G3 cell line using two different imaging platforms. First, for data of the WM989 NGFR-mNG2 cell line growing without drug, we used a Nikon Ti-E microscope encased in a plexiglass chamber ventilated with heated air and CO_2_. We took images at 60x magnification of mNG2 fluorescence every 6 hours and images of the iRFP nuclear reporter (H2B-iRFP670) every hour for 8.75 days. We chose these time intervals based on pilot experiments we performed to minimize overt signs of phototoxicity (cell death, growth inhibition, nuclear morphology changes) and enable the tracking of cell lineages. Second, for data in which we cultured these cells and then treated with vemurafenib, we used an IncuCyte S3 Live Cell Imaging Analysis System (Sartorius). We cultured the NGFR-mNG2 cell line on a 96-well plate inside the IncuCyte, which is fully contained within an incubator for long-term culture and time-lapse microscopy. With this system, we acquired images in green, red, and brighfield using a 10X objective at intervals of 2 hours over a total of 14.8 days. We added 500nM of vemurafenib after 5 days and 4 hours in culture, and then changed the media with vemurafenib every 3 days. We used these two different imaging platforms for their distinct advantages. The high magnification Nikon system allowed for the most accurate quantification of the NeonGreen2 signal, allowing us to measure the length of time of the NGFR fluctuations. Meanwhile, the IncuCyte is a more stable environment for time-lapse imaging that therefore induced less stress on the cells, and thus we used this platform for the longer experiments, particularly involving the additional stress of drug treatment.

### Time-lapse analysis

For tracking cell lineages and quantifying fluorescence signal of the NGFR-mNG2 WM989-A6-G3 cell line, we developed a set of publicly available tools for tracking cells in time-lapse images (https://bitbucket.org/arjunrajlaboratory/timelapsegui/wiki/Home). First, this pipeline uses an automated algorithm for nuclear segmentation to identify all the cells in each image. We then used a combination of automatic assignment of parents along with human-supervised annotation to fix errors to obtain lineage information. Next, we quantify the fluorescence signal from the NGFR-mNG2 by using the nuclear segment from each cell and calculating mean and median fluorescence intensity across these segments (Supp. Fig. 4). Of note, for the timelapses of AXL-high and EGFR-high sorted cells that lacked the H2B-iRFP670 nuclear marker, we had to manually mark all the cells that we wanted to analyze because the lack of nuclear markers precluded automatic segmentation. This same method of analysis was applied to all data acquired on the IncuCyte platform.. Our subsequent analysis consisted of building a custom data structure in MATLAB to contain each lineage and a series of plotting functions to allow us to plot the fluorescence intensity from an entire lineage (or part of a lineage) over the length of these experiments. For the lineages derived from AXL-high and EGFR-high sorted cells, the final frames were manually registered to images acquired after repeated immunofluorescence staining of live cells. The code for all the downstream processing is available on the Dropbox here: https://www.dropbox.com/sh/q2kbatojibljg8j/AABiFvNvsQm288tC3n-4qn5Za?dl=0 and https://www.dropbox.com/sh/zrc0g9vtxewcctl/AABVYCaXHYyvWFPOx4ipXLJGa?dl=0.

### RNA FISH

We designed custom oligonucleotide probe sets complementary to our genes of interest using custom probe design software written in MATLAB (code freely available for non-commercial use here https://flintbox.com/public/project/50547/) and ordered them with an amine group on the 3’ end (probe sequences available in Supp. Table 2). We pooled 15-32 oligonucleotides targeting each gene and coupled each set of probes to either Cy3 (GE Healthcare), Alexa 594 (Life Technologies), Atto647N or Atto 700 (Atto-Tec). We performed single-molecule RNA FISH as described in Raj *et al.* (2008) for multiple cycles of hybridization. We fixed cells in 4% formaldehyde solution for 10 minutes at room temperature, permeabilized in 70% ethanol, and stored samples at 4C. For hybridization, we first washed samples with washing buffer containing 10% formamide and 2x SSC. We then applied hybridization buffer containing custom RNA FISH probes and 10% formamide, 2x SSC, and 10% dextran sulfate. We hybridized samples overnight at 37°C and then performed two cycles of 30 minute washes at 37°C with washing buffer. For imaging, we first DAPI stained the cells and then transferred them to 2x SSC. The sequences of the RNA FISH probes used in this manuscript are located here: https://www.dropbox.com/s/dplhgrvznny1hau/supplement_probe_sequences.xlsx?dl=0

### RNA FISH imaging

We imaged RNA FISH samples on an inverted Nikon TI-E microscope with a 60x Plan-Apo or a 100x Plan-Apo using filter sets for DAPI, Cy3, Atto647N, Alexa594, and Atto700. We took images in either z-stacks of 30 planes at 0.3µm intervals using custom journals built in Metamorph or tiled grids of single-plane images using Metamorph Scan Slide Application. We used a Nikon Perfect Focus system to maintain focus across the imaging area.

### RNA FISH image analysis

For analysis of gridded image scans, we used custom MATLAB software designed for the analysis in (Shaffer et al., 2017). This pipeline consists of first segmenting the nuclei of individual cells using DAPI images. Next, the software calculates regional maxima for all RNA FISH dyes and then the user specifies a global threshold for calling individual spots. Through a GUI interface the user then reviews the high expressing cells and uses editing tools to remove artifacts or autofluorescent debris. Lastly, we constructed tables containing RNA FISH spot counts for each gene in individual cells.

For image z-stacks, we used custom MATLAB software to count spots for each cell. Briefly, this image analysis pipeline includes manual segmentation of cell boundaries, thresholding of each channel in each cell to identify individual spots, and then extraction of spot counts for each gene in each cell. The software for analysis of RNA FISH images is available on bitbucket here https://bitbucket.org/arjunrajlaboratory/rajlabimagetools/wiki/Home. After extracting spot counts from either data format, we performed the remainder of the analysis of mRNA distributions in R. We calculated Gini coefficients (as described in Ref. (Jiang et al., 2016)) for each gene using the “ineq” package. The code for this analysis is on the Dropbox here https://www.dropbox.com/s/n8cppr17b3bgssf/gini_coef_analysis.R?dl=0.

### Spatial analysis

We used spatial single cell analysis to enable us to independently measure the heritability of high expression states. We sparsely plated cells (WM989-A6, MDA-MB-231-D4, WM983B) on a 2-well chambered coverglass, and then we allowed the cells to grow for 10 days (sometimes fewer days for MDA-MB-231-D4 if the cells grew faster), at which point we performed iterative RNA FISH, image analysis, and thresholding for high expression as described above. Intuitively, the stronger the heritability (i.e., over several generations), the larger the clusters of high-expressing cells we would find. To quantify clustering, we created, for each cell in the dataset, a “bucket” consisting of the 20 (or 50, 100, 200) closest cells and then kept track of the number of high-expressing cells in that bucket. We then computed the variance and the mean of this number across all buckets, allowing us to then calculate the heritability index, which is the Fano factor (defined as the variance divided by the mean). In the case of complete spatial randomness, the distribution would be Poisson, and the heritability index would be 1. To verify this null distribution, we permuted the label of cells as jackpots or non-jackpots uniformly at random 1000 times, and recomputed the heritability index for each permutation. This approach allowed us to compute 95% confidence intervals for the null distribution given our particular spatial configuration of cells; the data for null distributions is not shown, but is available online here: https://www.dropbox.com/sh/m1rdxjrfeo6be3m/AADphkUBGT0zgppppssP_LbSa?dl=0.

### Network community identification

For each pair of significantly heritable genes (from gene lists described in the MemorySeq RNA sequencing analysis section of the Methods), we calculated the Pearson correlation coefficient between their expression across clones. This procedure resulted in a symmetric weighted matrix of size 227 genes x 227 genes in WM989 (and 230 genes x 230 genes in WM983B, as well as MDA-MB-231-D4). We represent these matrices as undirected weighted networks with nodes corresponding to genes, and with the weight of edges between nodes corresponding to the value of the correlation coefficient. Within this network, we sought tightly connected groups of nodes within this network known as *network communities*. We performed *k*-clique community detection (Derényi et al., 2005; Jonsson et al., 2006; Palla et al., 2005) with k=4 on binarized networks created by thresholding the original weighted network at decreasing values (Giusti et al., 2015; Rieck et al., 2018). More specifically, a *k*-clique is a collection of *k* nodes that are all-to-all connected and a *k-*clique community is a collection of *k*-cliques that are connected through adjacent *k*-cliques (two *k-*cliques are adjacent if they share *k-1* nodes). Repeatedly thresholding the network at decreasing values of the edge weight creates a sequence of binary graphs, each of which is included in the next. Since the addition of edges to a binary graph can only enlarge or merge *k*-clique communities, we can track communities from one binary network to the next in a well-defined manner. This mapping allows us to both observe which nodes were included in the community at slightly lower threshold values and to qualitatively assign statistical significance to communities based on the range of threshold values for which they stay isolated from the rest of the network.

## Acknowledgements

Thank Siyu Feng, David Brown, and Bo Huang for constructs pSFFV_mNG2(1-10), and Christian Meyer for discussions about sort-relaxation experiments. We thank Caroline Bartman, Yogesh Goyal, and other members of the Raj lab for many useful suggestions. We thank the Flow Cytometry Core Laboratory at the Children’s Hospital of Philadelphia Research Institute for assistance in designing and performing FACS. SMS acknowledges support from NIH F30 AI114475. BLE acknowledges support from NIH training grants T32GM007170 and T32HG000046. RRH acknowledges support from the NSF Graduate Research Fellowship DGE-1845298. AS acknowledges support by NIH grants 5R01GM124446 and 5R01GM126557. AES and DSB acknowledge support from the John D. and Catherine T. MacArthur Foundation, the Alfred P. Sloan Foundation, and the ISI Foundation. AR acknowledges NIH/NCI PSOC award number U54 CA193417, NSF CAREER 1350601, P30 CA016520, SPORE P50 CA174523, NIH U01 CA227550, NIH 4DN U01 HL129998, NIH Center for Photogenomics (RM1 HG007743), and the Tara Miller Foundation.

