## Supplementary Figures for "Memory sequencing reveals heritable single cell gene expression programs associated with distinct cellular behaviors"

### Supplementary Figure 1

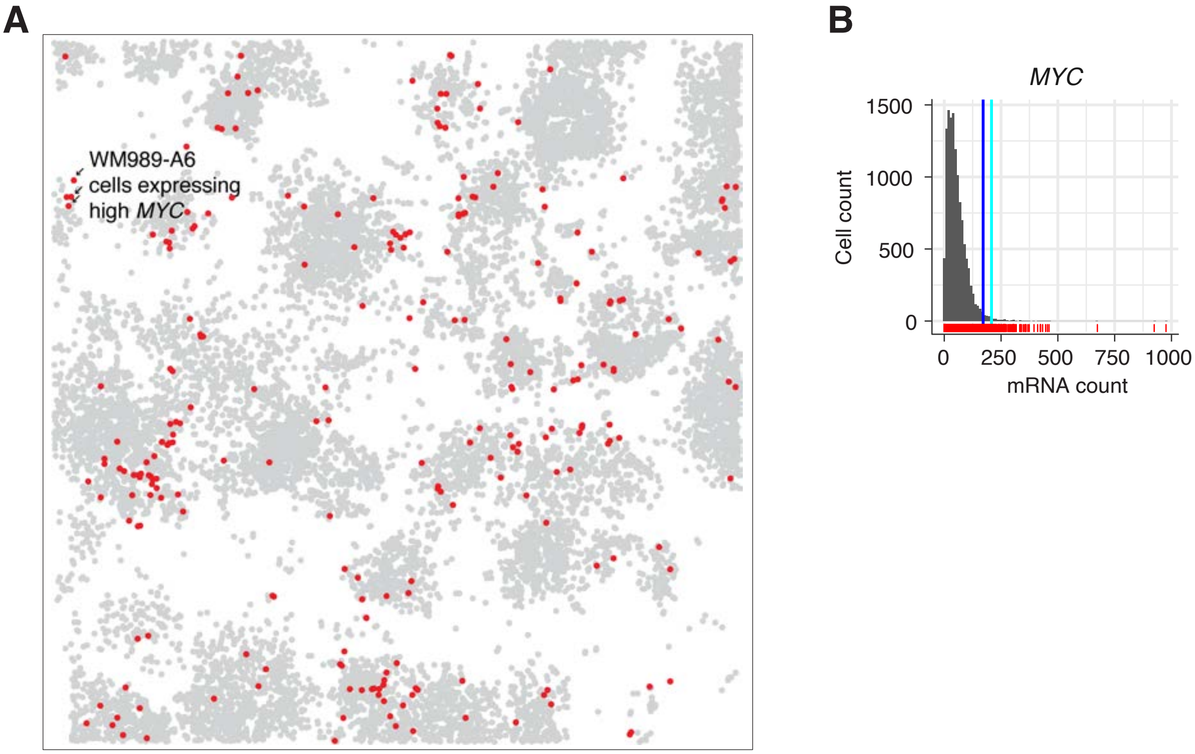


**Supplementary Figure 1. MYC expresses in a highly variable but not heritable pattern.** We measured expression of MYC in WM989-A6 cells grown on culture dishes for 10 days. **A.** Spatial position of each cell in culture with the top 2% of *MYC* expressing cells in red and the rest of cells labeled in gray. The degree of spatial clustering was minimal, reflecting the low heritability of the high MYC expression level cellular state. **B.** Histogram of *MYC* mRNA levels across individual cells, showing the large amount of variability in MYC expression. The rightmost line (light blue) marks the top 1% of cells, the left line (dark blue) marks the top 2%.

#### Supplementary Figure 2

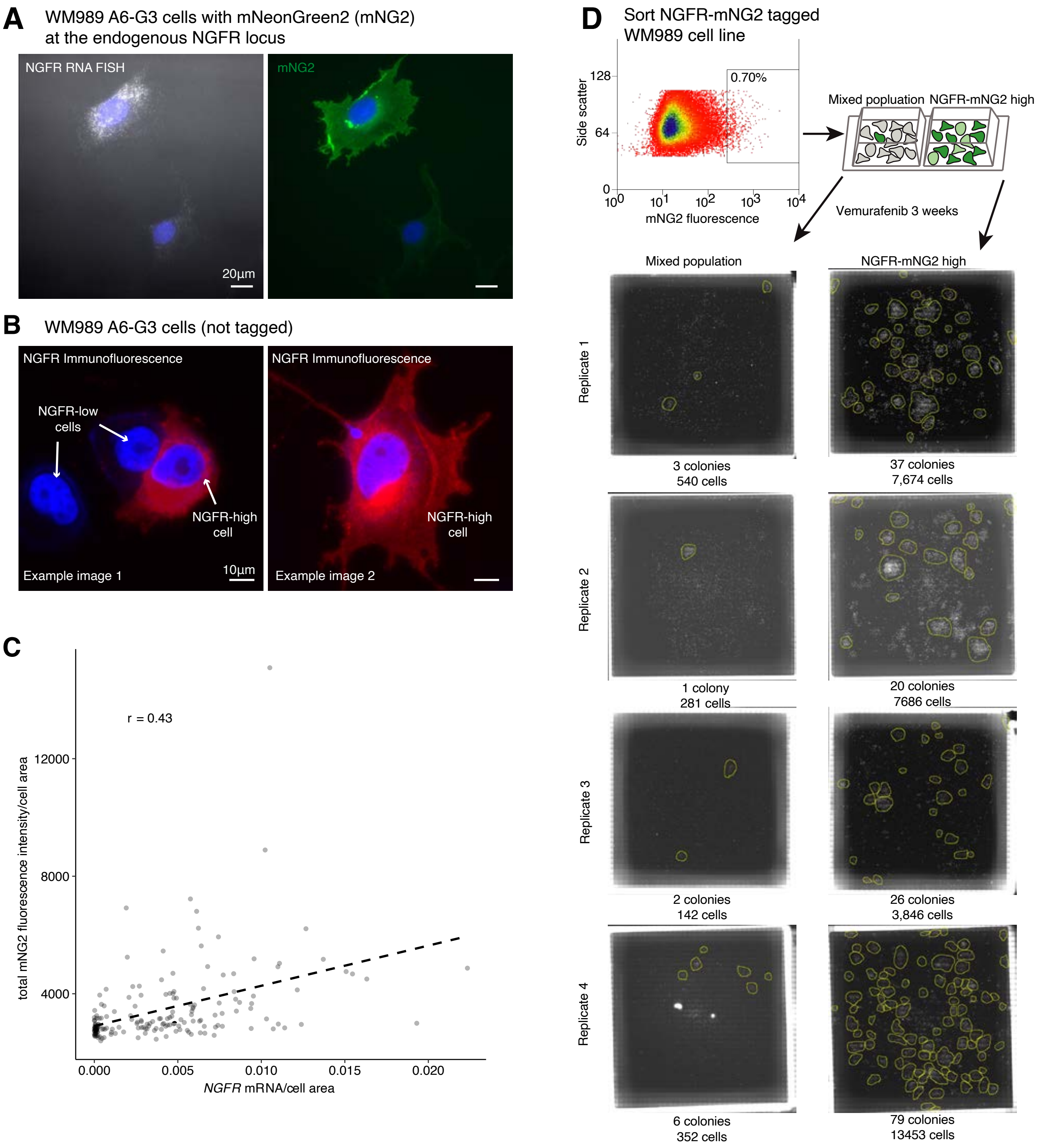


**Supplementary Figure 2. Validation of mNG2 fluorescence as a marker of *NGFR* expression**. We sorted NGFR-mNG2 tagged WM989-A6-G3 cells then performed single-molecule RNA FISH to measure NGFR mRNA levels. **A.** Representative max merged image of NGFR-mNG2 cell line. Scale bars are 20µm. **B.** Immunostaining of endogenous NGFR in untagged WM989-A6-G3 shows a similar fluorescence localization pattern as NGFR-mNG2. Scale bars are 10µm. **C.** Scatterplot of data from 188 analyzed cells show that mNG2 fluorescence per cell area correlates with NGFR mRNA per cell area in single cells. **D.** We sorted bulk and mNG2-high (top 0.5-1%) WM989-A6-G3 then treated cells for 3 weeks with 1µM vemurafenib. After fixation and staining nuclei with DAPI, we imaged plates and quantified drug resistant colonies using publicly available software (see Methods). The indicated number of cells correspond to cells within colonies. As with antibody staining for NGFR, sorting NGFR-mNG2 tagged WM989 A6-G3 by mNG2 fluorescence enriches for vemurafenib resistant cells

### Supplementary Figure 3

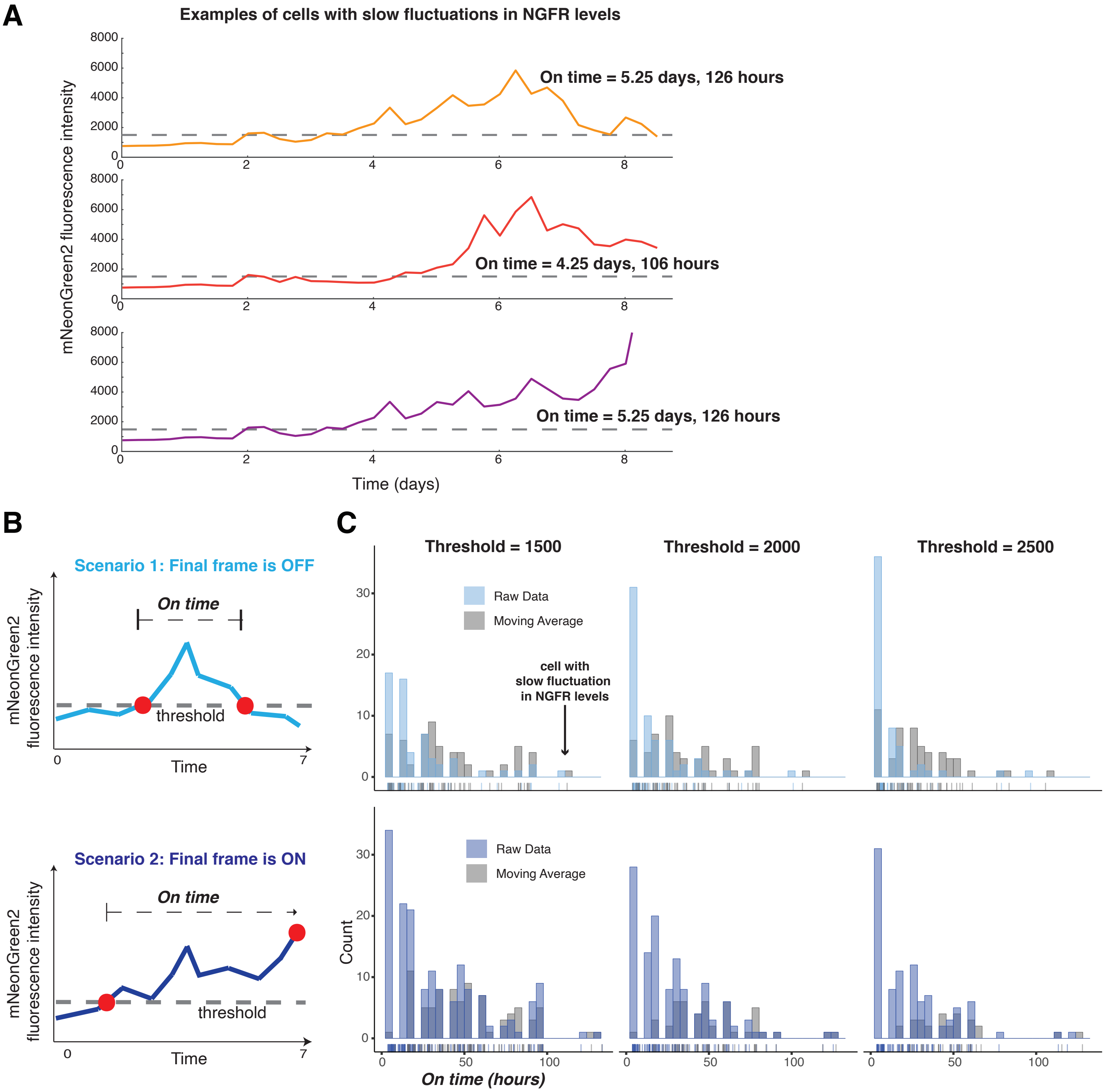


**Supplementary Figure 3. Fluctuations in NGFR levels can persist for at least 5.25 days.** We performed time-lapse imaging of WM989-A6-G3 cells with NGFR tagged with mNeonGreen2 at the endogenous locus. We tracked cells through these images and quantified the mNeonGreen2 fluorescence signal to measure the length of time that cells reside in the NGFR-high cell state. **A.** Plots of mNeonGreen2 fluorescence intensity over time for three example cells that enter the NGFR-high state and remain high for 4.25 to 5.25 days, thereby demonstrating memory. These cells are from a lineage for which we manually curated data through 8.75 days. **B.** Cartoons depicting the two outcomes for NGFR-high cells. In scenario 1, cells pass the threshold and remain NGFR-high for a period of time (*on time*) and then fall below the threshold and are NGFR-low. In scenario 2, the cells become NGFR-high, but do not fall below the threshold during the length of this experiment (7 days). Given that these two scenarios are distinct we analyzed them separately. **C.** Histograms of the *on time* for all cells through 7 days of imaging (shorter data set than A, as we manually reviewed all cells up to this time point). We considered a range of thresholds (1500, 2000, 2500) for determining when individual cells are in the NGFR-high cell state. The blue (both dark and light blue) show the raw data for the *on time*, while the gray shows the on times that result from using a moving average of the time trace using a 5 point median filter. At a threshold of 1500, the average *on time* is 40 hours for scenario 1 (light blue) and 52 hours for scenario 2 (dark blue).

### Supplementary Figure 4

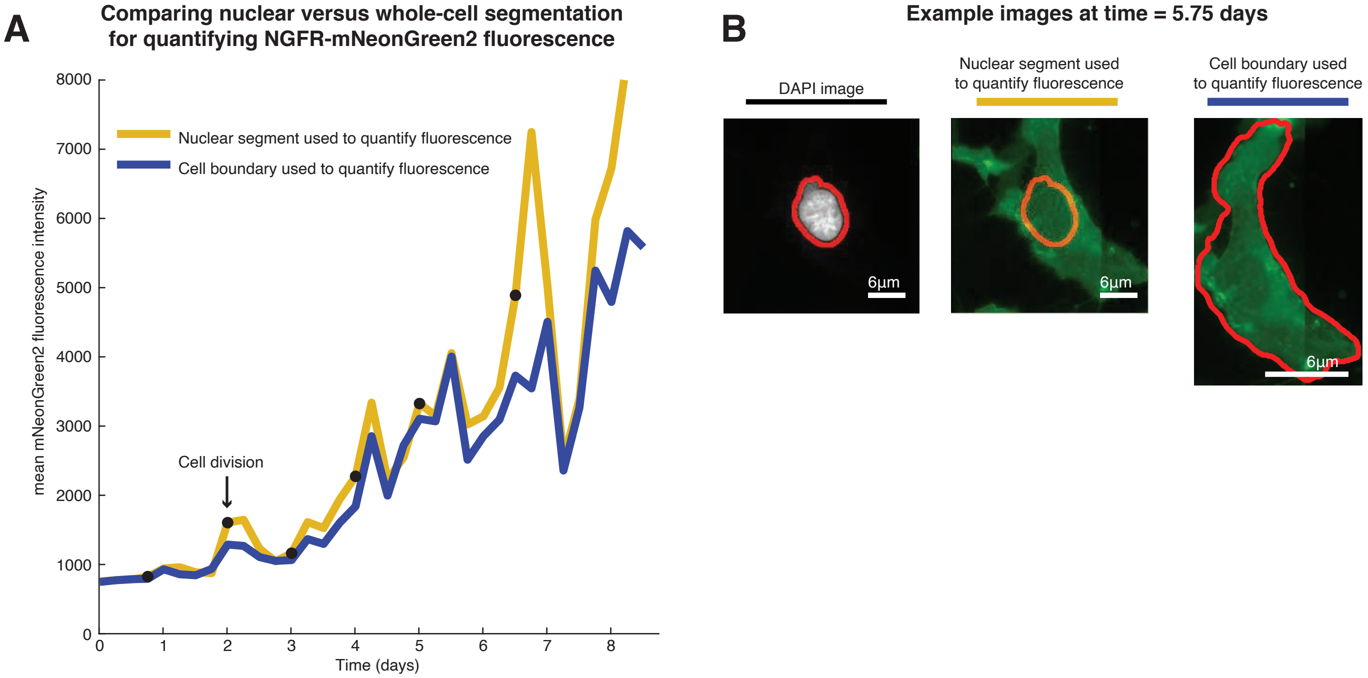


**Supplementary Figure 4. Comparison of fluorescence quantification for NGFR-mNG2 using nuclear and whole-cell segmentation.** We developed a WM989 cell line with mNeonGreen2 at the endogenous NGFR locus. We performed time-lapse imaging of this cell line over 8.75 days. For analysis of this data, we wanted to determine if the nucleus area could be used for quantifying the mean fluorescence intensity from the NGFR mNG2 tagged cell line. We tracked one cell through 8.75 days and performed manual segmentation of the whole-cell at each time point. We then compared the mean fluorescence intensity from our manual segment to the mean fluorescence intensity observed over the nucleus as determined by an automatic segmentation algorithm (described in the methods). We found that these two approaches gave similar fluorescence intensity patterns over the 8.75 days. **A.** This plot shows the mean fluorescence intensity for mNeonGreen2 over time using the two different approaches. The blue line is the manual segmentation of the whole-cell and the yellow line is the automatic nuclear segmentation. **B.** Example images from each segmentation technique at time point 5.75 days. The scale bars are 6 µm.

### Supplementary Figure 5

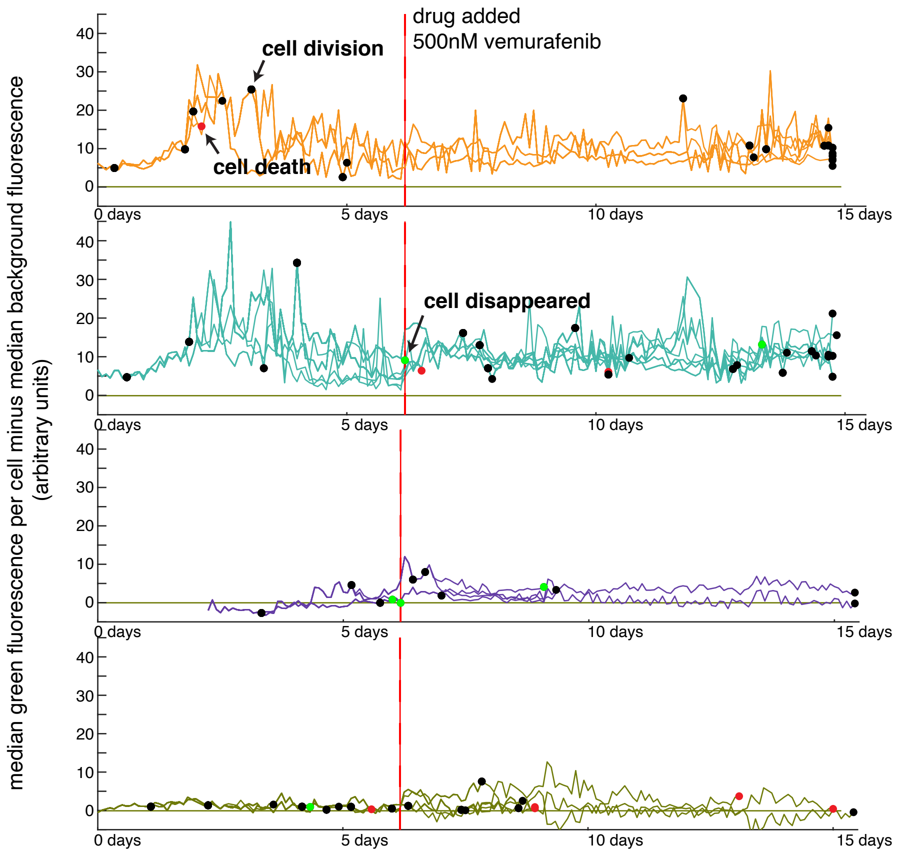


**Supplementary Figure 5. NGFR-high cells survive vemurafenib treatment and continue to proliferate more than NGFR-low cells.** We performed time-lapse imaging of WM989-A6-G3 cells in which the endogenous NGFR is tagged with mNeonGreen2. We acquired images every 2 hours for 14.8 days and added 500nM of vemurafenib starting after 6 days and 4 hours. Our analysis consisted of tracking lineages and then quantifying the green fluorescence signal from the NGFR-mNeonGreen2. This plot shows the fluorescence signal over time for four lineages in this data set. The y-axis is the median fluorescence over each cell segment minus the median fluorescence of the background for the image. The black dots are cell division, the red dots are cell death, and the green dots are when a cell disappears from the culture dish. The red line marks the time point at which we added 500nM vemurafenib to the sample.

### Supplementary Figure 6

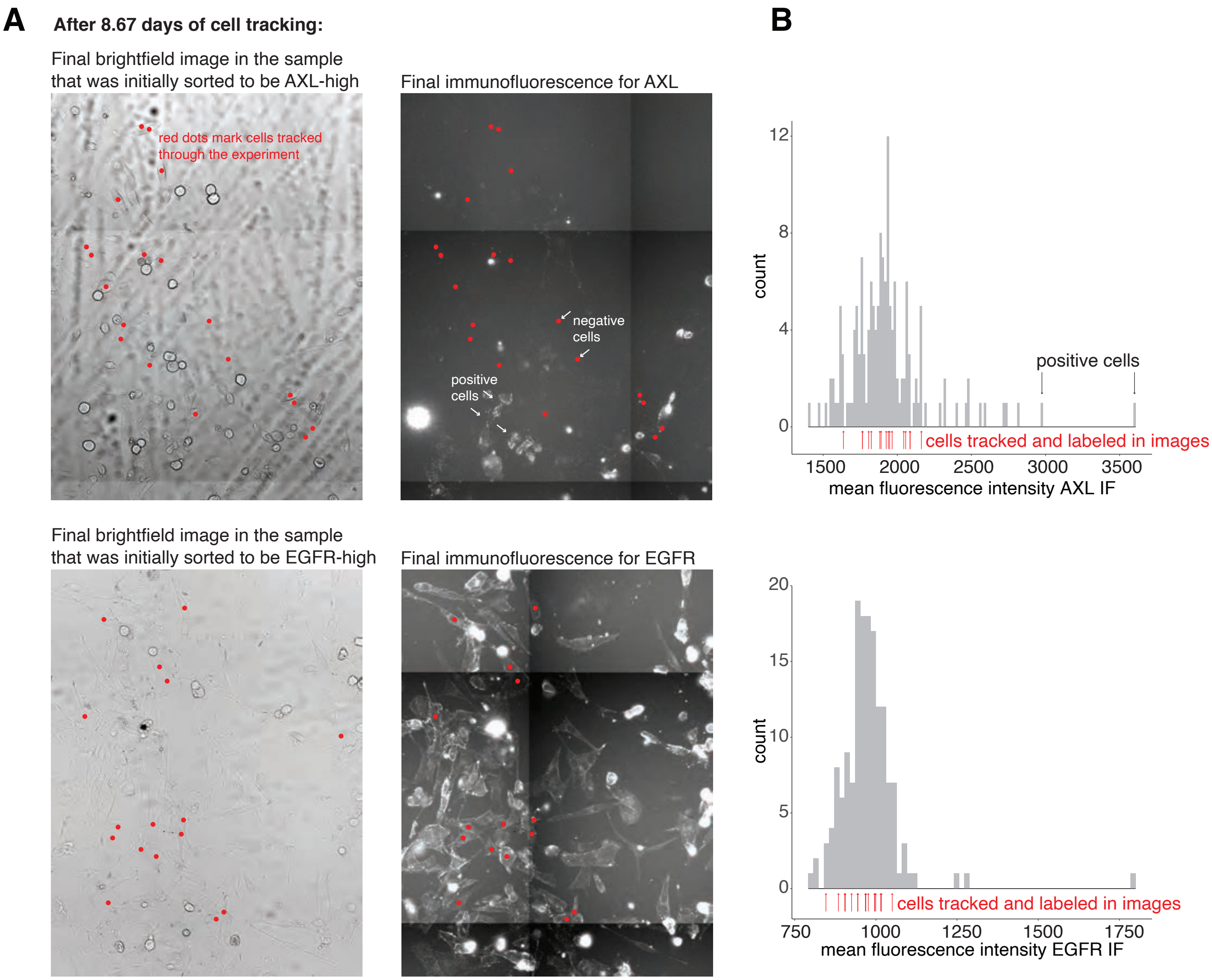


**Supplementary Figure 6. Immunofluorescence after tracking of cells based on transmitted light imaging reveals emergent heterogeneity.** A. Fields of view taken after 8.67 days of imaging in both transmitted light and immunofluorescence targeting either AXL or EGFR. All tracked cells are labeled with red points. B. Histograms of number of average immunofluorescence signal intensity of all cells after the imaging period. Red arrows mark the intensity values for the cells labeled by dots in the images which were tracked through the 8.67 days. All scale bars are 10 $\mu$m long.

#### Supplementary Figure 7

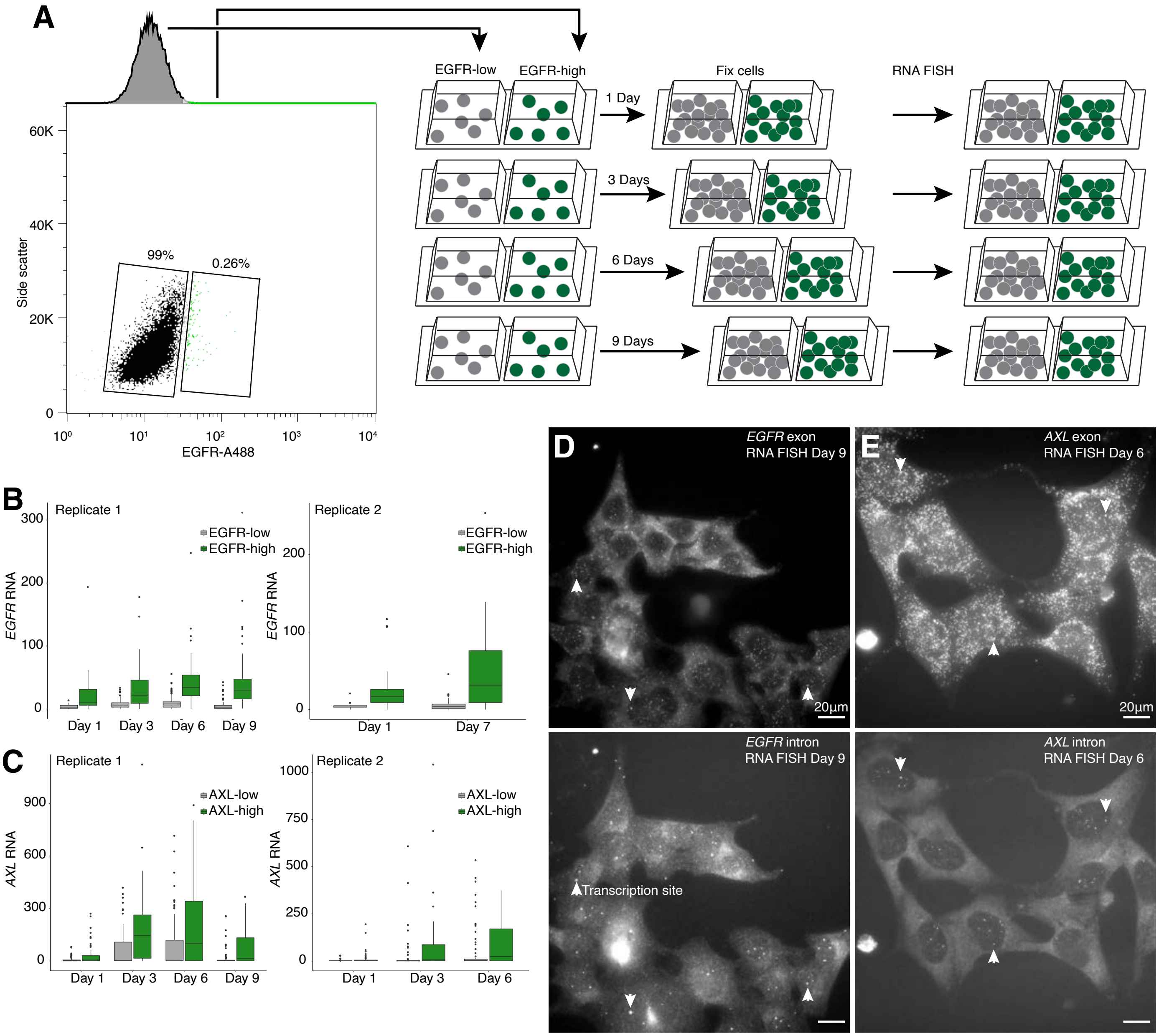


**Supplementary Figure 7. FACS analysis shows memory over 5-9 days: A**. To estimate the transcriptional memory of *EGFR* and *AXL* in WM989 A6-G3, we sorted out high (top ~ 0.3% for EGFR and top ~ 0.5% for AXL) and low expressing cells then cultured these cells for up to 9 days before fixing and performing RNA FISH. **B-C**. We found persistence of elevated *EGFR* and *AXL* expression for up to 9 days in EGFR-high and AXL-high sorted cells, respectively. For *AXL*, we saw an initial increase in AXL expression after sorting followed by a decrease towards baseline at Day 9, consistent with a long-lived but ultimately transient gene-expression state. The initial increase in *AXL* expression may be due to paracrine signaling or stress from sorting. **D-E**. Representative images showing clusters of EGFR-high and AXL-high expressing cells. Arrowhead point towards examples of transcription sites marked by colocalization of exon-targeting and intron-targeting RNA FISH probes. The presence of multiple transcription sites in these cells suggested that the observed gene-expression memory is due to persistence of active transcription rather than slower division or RNA degradation rates.

#### Supplementary Figure 8

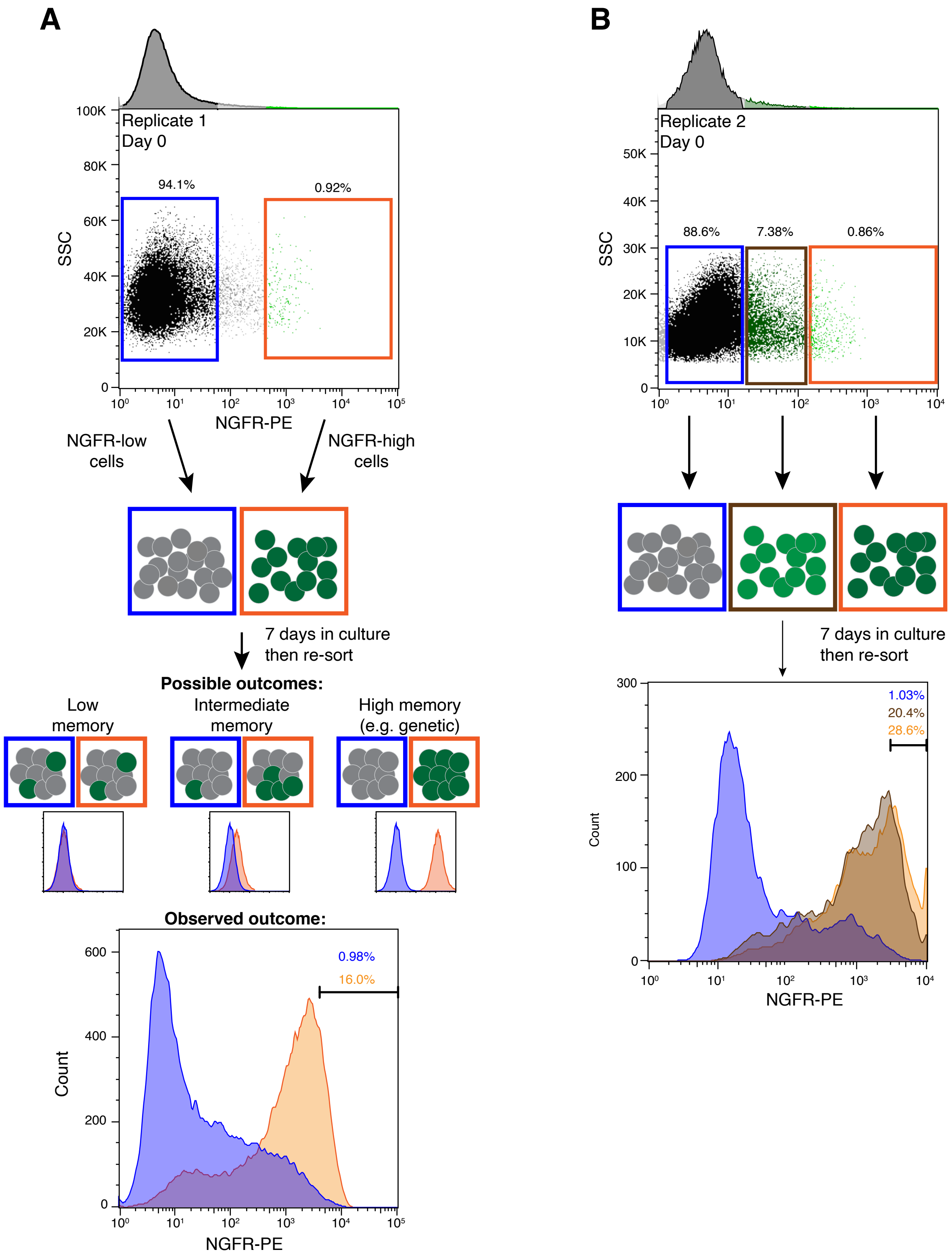


**Supplementary Figure 8. Isolated NGFR-high WM989 A6-G3 cells revert towards baseline NGFR levels over the course of 7 days in culture. A-B**. We labeled W989 A6-G3 cells with a Phycoerythrin-Cyanine7(PE-Cy7) conjugated antibody targeting NGFR, then sorted equal numbers of cells from the indicated gates. After 7 days in culture, we re-stained the cells and measured fluorescence intensity by flow cytometry. At day 7, the distribution of NGFR-PE-Cy7 intensities is shifted higher for the NGFR-high samples relative to the NGFR-low samples, although the distributions are closer than at day 0.

### Supplementary Figure 9

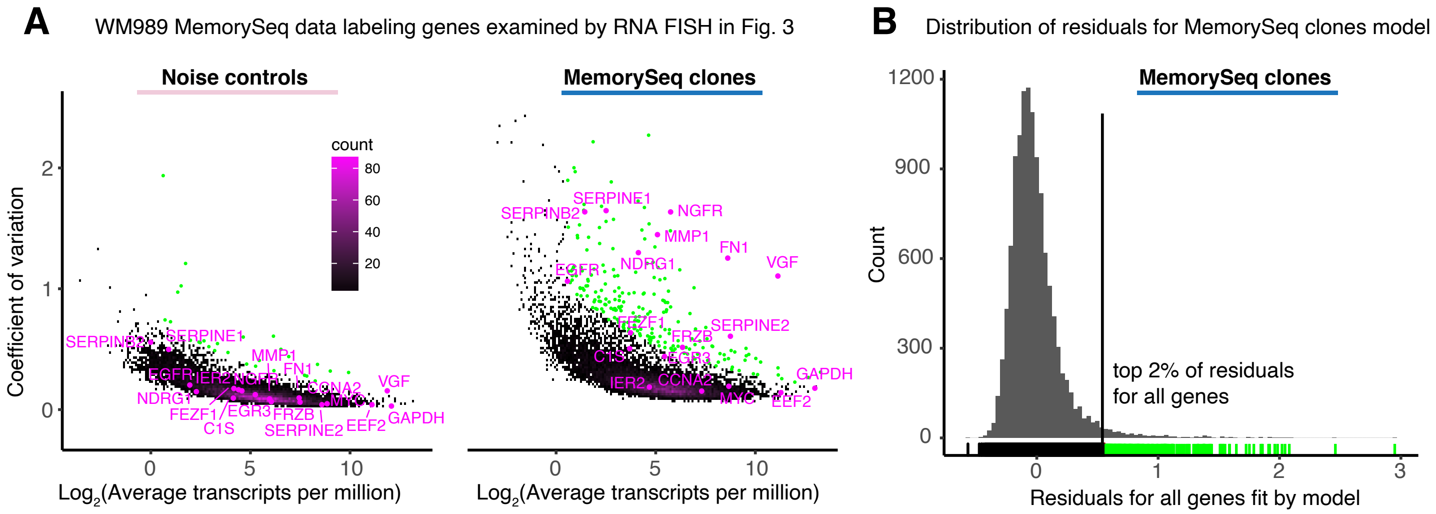


**Supplementary Figure 9. MemorySeq data for WM989-A6 melanoma cells with labels showing the genes selected for RNA FISH in Fig. 3. A.** Plot of coefficient of variation versus the log2 mean expression for all genes. These plots are similar to the plots in Fig. 1D, expect the genes labeled in pink are the genes selected for RNA FISH in Fig. 3. The points labeled with the green dots are genes that passed the threshold for selection as a heritable gene. **B.** Histogram showing the overall distribution for the residuals for all genes fitted by the model on the MemorySeq clones. To select heritable genes, we used a cutoff at the top 2% of the residuals. This cutoff is highlighted in green on the tick marks below the histogram.

### Supplementary Figure 10

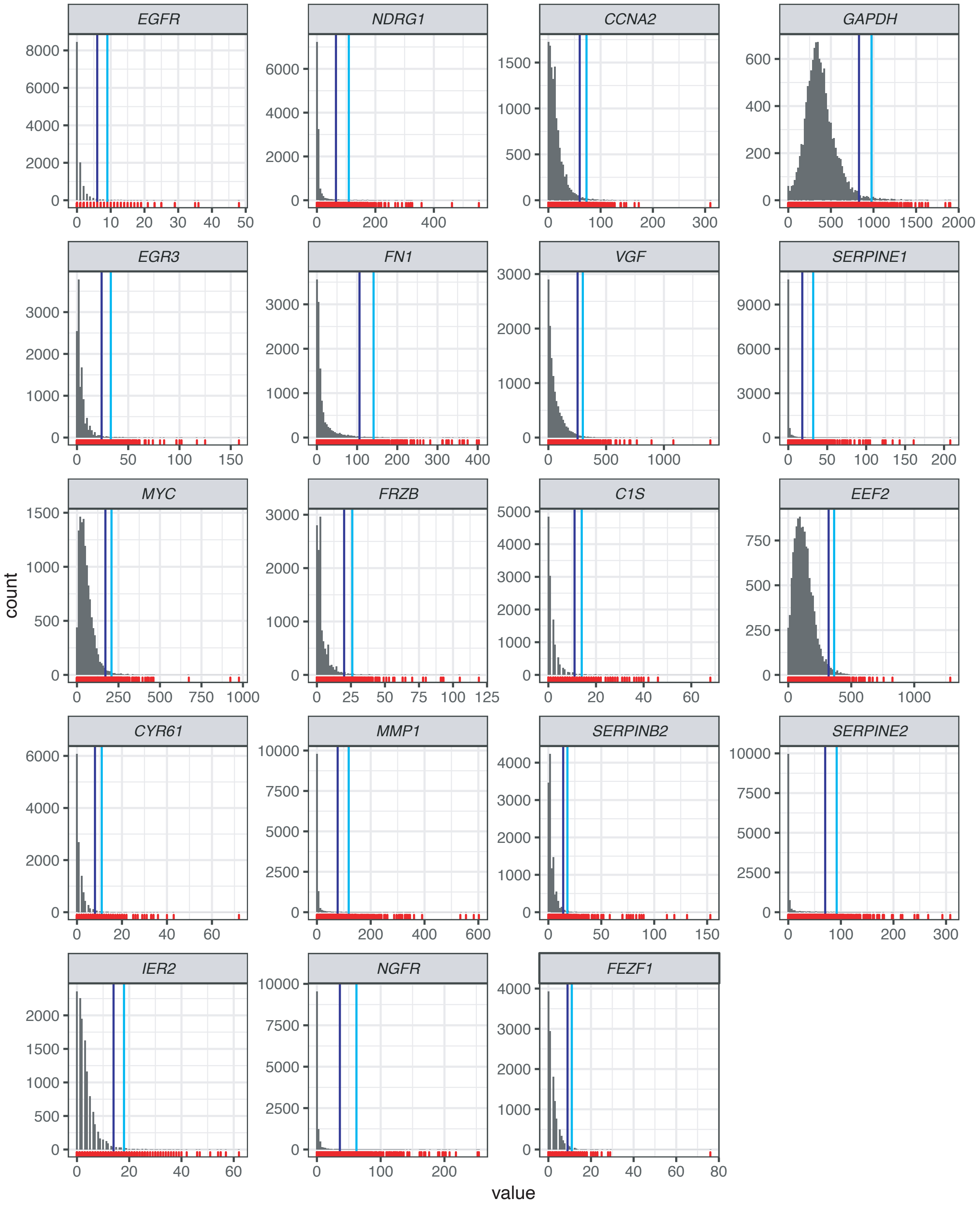


**Supplementary Figure 10. Histograms showing the distribution of expression levels in single cells as measured by RNA FISH.** We plated WM989 cells sparsely on a dish and then allowed them to grow into clusters over 10 number of days. We performed RNA FISH for a panel of 19 genes and then quantified expression for each gene in single cells. The histograms show these expression levels for 12,192 cells total. One of two replicates is shown. Equivalent plots for replicates and other cell lines are available on the paper dropbox.

### Supplementary Figure 11

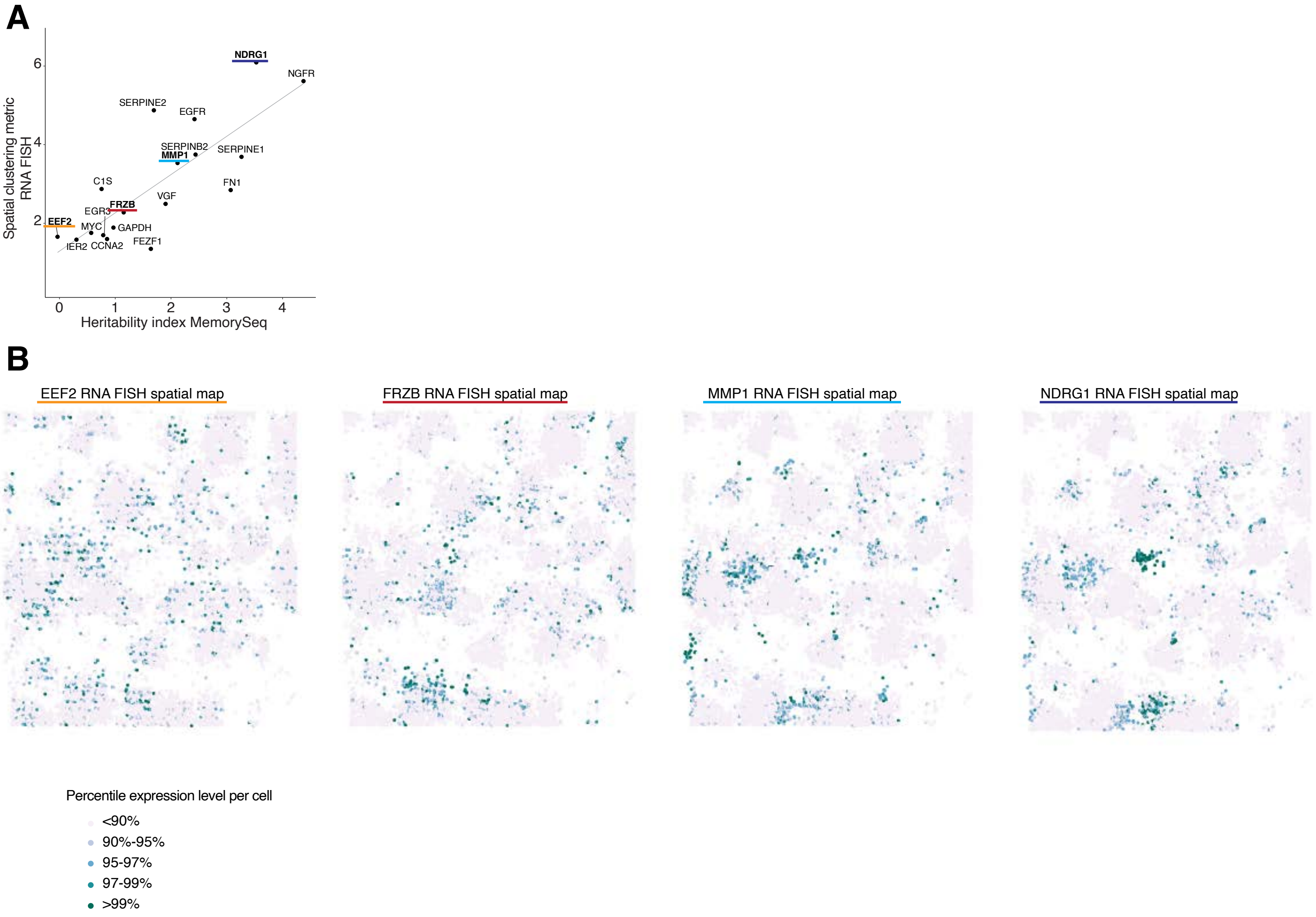


**Supplementary Figure 11. Spatial maps for the data shown in Fig. 3A-D with each cell color coded by the percentile of number of RNAs. A.** Same plot as shown in Fig. 3C with genes underlined that are displayed in B. **B.** Spatial maps of the x and y position of cells with color coding to label different percentile cutoffs for determining if a cell has “high” expression. For these experiments, we plated cells sparsely on the dish and then allowed them to grow into clusters for 10 days. Here, we expect sister cells and other related cells to be near each other on the dish. We then fixed these samples and performed RNA FISH for a panel of 19 genes. We used spatial position as a proxy for cell relatedness and looked for clustering of the cells expressing these genes to indicate memory of the transcriptional “high” state. These plots show 12,192 cells total. The color indicates the percentile cut-off for determining which cells have “high” expression.

### Supplementary Figure 12

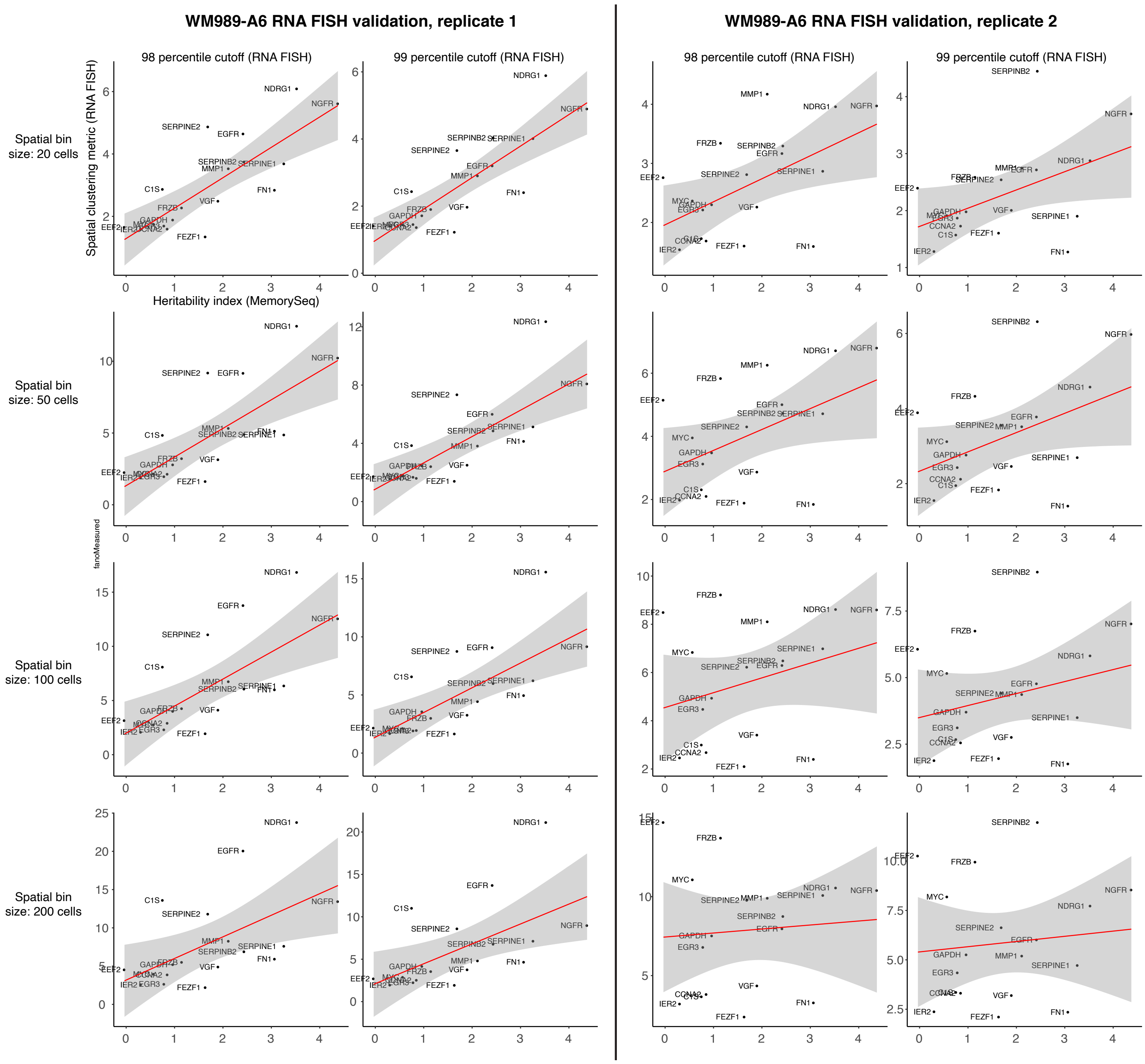


**Supplementary Figure 12. Spatial single cell RNA FISH analysis validation of MemorySeq.** We performed analysis of single molecule RNA FISH for the indicated genes as per Fig. 3. Shown are two biological replicates (left and right). Within each replicate, we analyzed different cutoffs for cells to be considered to be in the high expression level state (either the top 1 percent of cells or the top 2 percent of cells). We also increased the bin size for each neighborhood for analysis, which is quantified as the number of neighboring cells included in each bin. We quantified the skewness across MemorySeq clones vs. the Spatial clustering metric, which is the Fano factor (variance/mean) in the number of positive cells per bin across all bins of indicated size. We found the correspondence decreased once the bin size reached 100, and that using the top 1 percent of cells also led to a tighter correlation. The decreased correlation at higher bin size likely reflects the fact that at larger bin sizes, more clusters would merge together, thus decreasing apparent Fano factor across bins. (To see this, take the limiting case of just two bins, in which the variance between the two would be relatively minimal.)

### Supplementary Figure 13

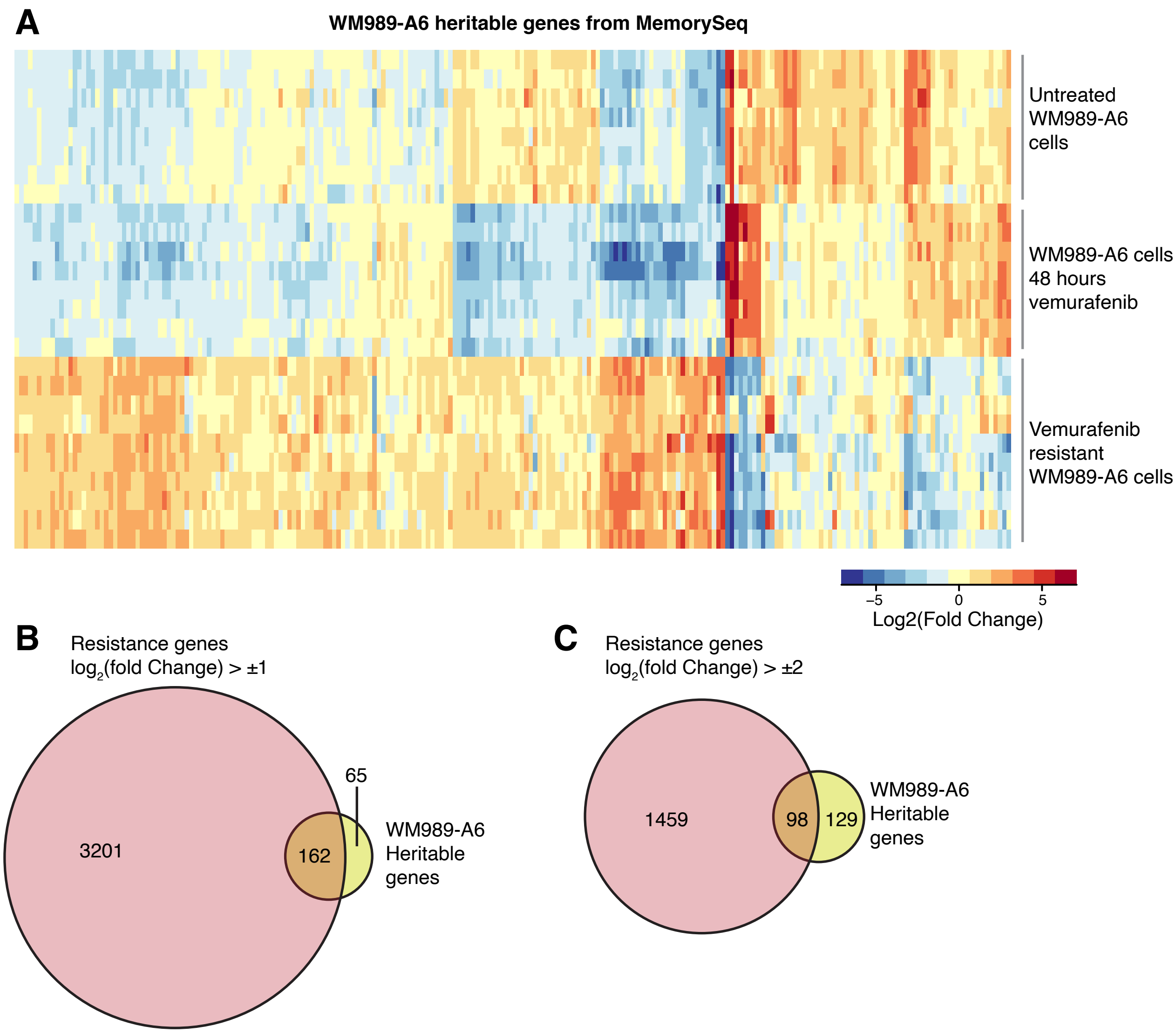


**Supplementary Figure 13. Genes identified by MemorySeq and their expression levels in stably resistant melanoma cells.** We wondered how the expression levels of genes identified by MemorySeq changed during the process of acquiring resistance (WM989-A6 cells) using data from Shaffer *et al.* 2017. **A.** Each column is a gene identified by MemorySeq as being heritable. Each row is a biological replicate, from bulk untreated cells, cells treated with vemurafenib for 48 hours (to account for drug-response effects) and the stably resistant cell lines. **B, C.** Venn diagram showing which genes were associated with resistance together with being identified as heritable by MemorySeq. We ran the analysis at two different thresholds for fold change. Either way, we found that while a large fraction of heritable genes were also resistance-associated genes (over 70% for a two-fold change cutoff), only a very small proportion of total resistance genes were identified by MemorySeq as being heritable, in agreement with the conclusions of Shaffer *et al.* 2017.

### Supplementary Figure 14

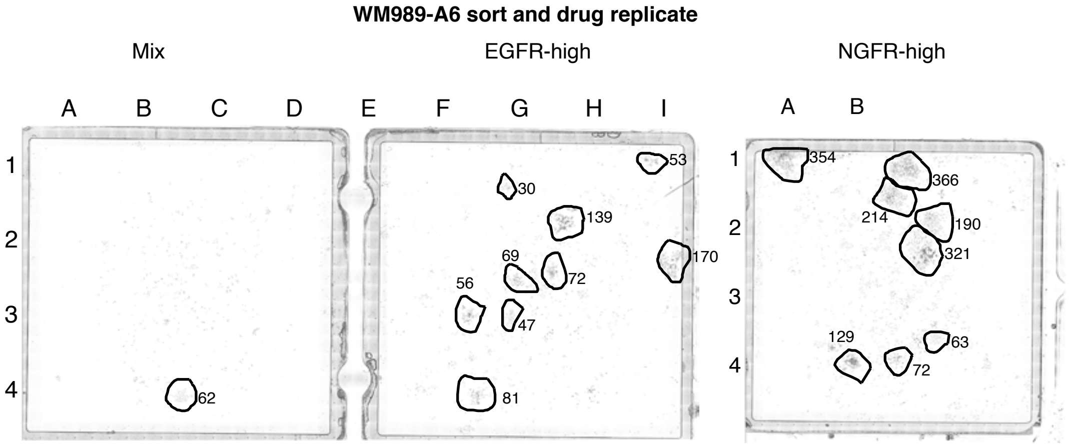


**Supplementary Figure 14. Replicate of NGFR sorted cells grown in trametinib.** Biological replicate of the data shown in Fig. 3. We used fluorescence activated cell sorting to isolate the NGFR-high and EGFR-high subpopulation of WM989-A6 cells, cultured for 8-16 hours and then subjected to trametinib treatment at 10nM (MEK inhibitor) for 3 weeks. Image shows number of resistant colonies (circled) along with number of cells within the resistant colony as indicated.

### Supplementary Figure 15

Overlapping heritable gene sets across WM989-A6, MDA-MB-231-D4, and WM983B-E9:

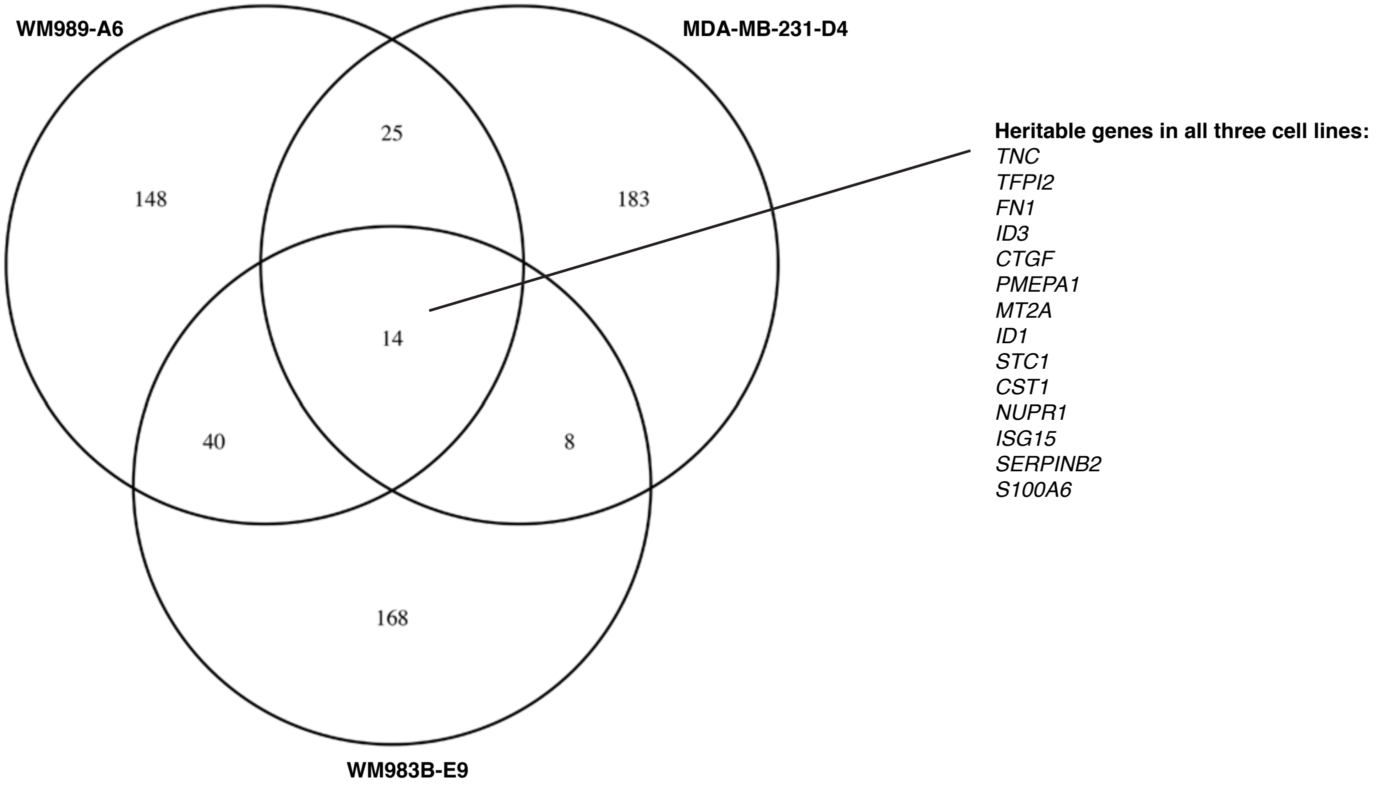


**Supplementary Figure 15. Overlap of MemorySeq hits across cell lines.** We wondered if genes identified by MemorySeq as heritable were the same across multiple cell lines. We found that most heritable genes were distinct between cell lines, perhaps reflecting the potential for cell-type specific heritable rare cell expression programs. There was a set of 14 genes (listed in figure) that were identified as heritable in all 3 cell lines.

### Supplementary Figure 16

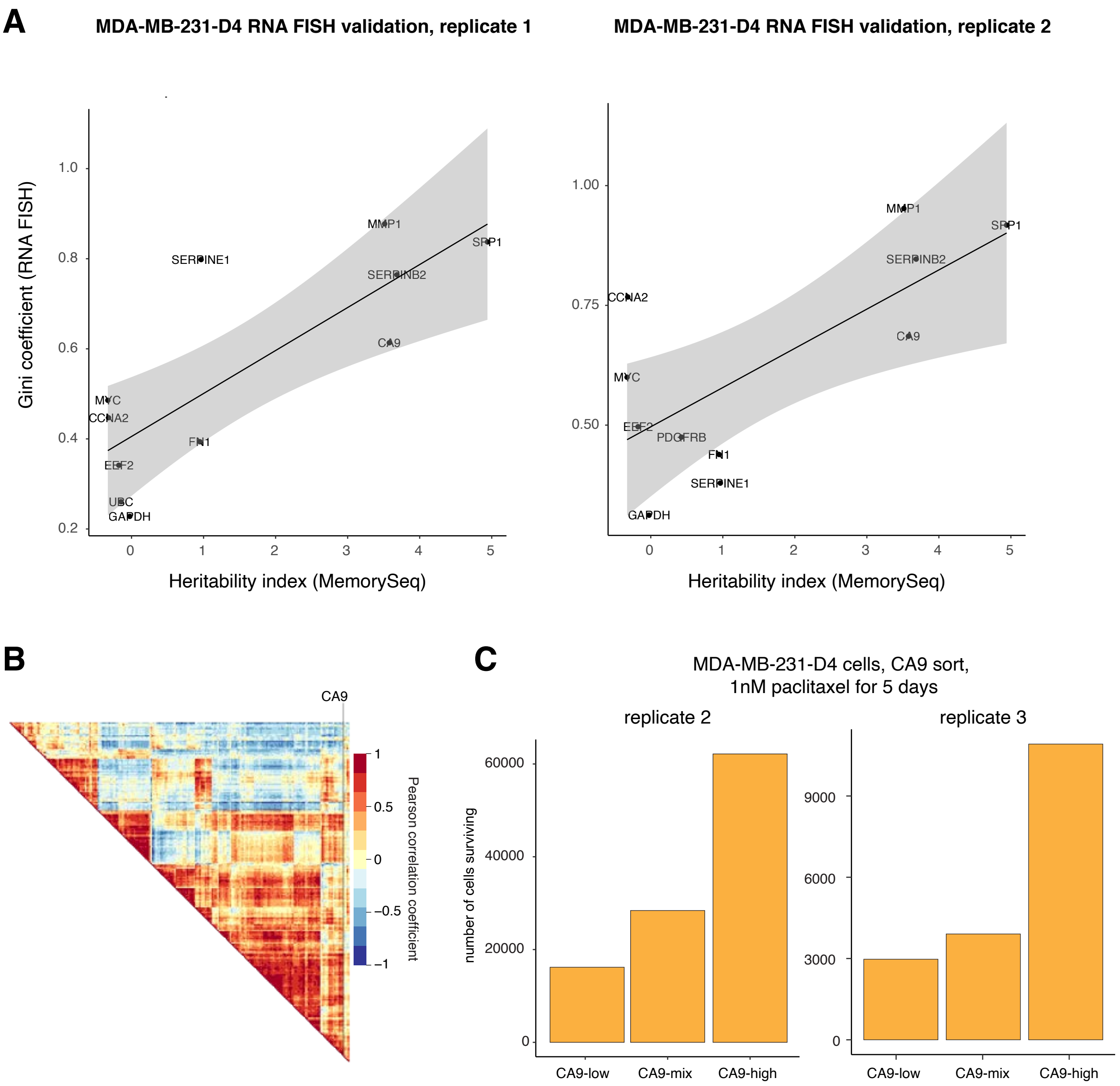


**Supplementary Figure 16. Comparison of heritability by MemorySeq to Gini coefficient for MDA-MB-231-D4 cells. A.** We plotted heritability index from MemorySeq (skewness across individual MemorySeq clones) vs. Gini coefficient as computed from single molecule RNA FISH across two biological replicates. Overall, we found a general correspondence between the two variables, as was seen for WM989-A6 cells, suggesting that MemorySeq also identifies genes that express in rare cells. **B.** Correlation heatmap for all pairs of heritable genes in MDA-MB-231-D4 MemorySeq. **C.** Replicate data for Fig. 4D. We stained cells with antibody targeting the CA9 surface marker and then sorted out the top 2-4% of cells, the lowest 2-4% of cells, and the total "mix" population into chamber wells, after which we applied paclitaxel 1 day after sorting for 5 days. Plots show the number of cells in each condition.

### Supplementary Figure 17

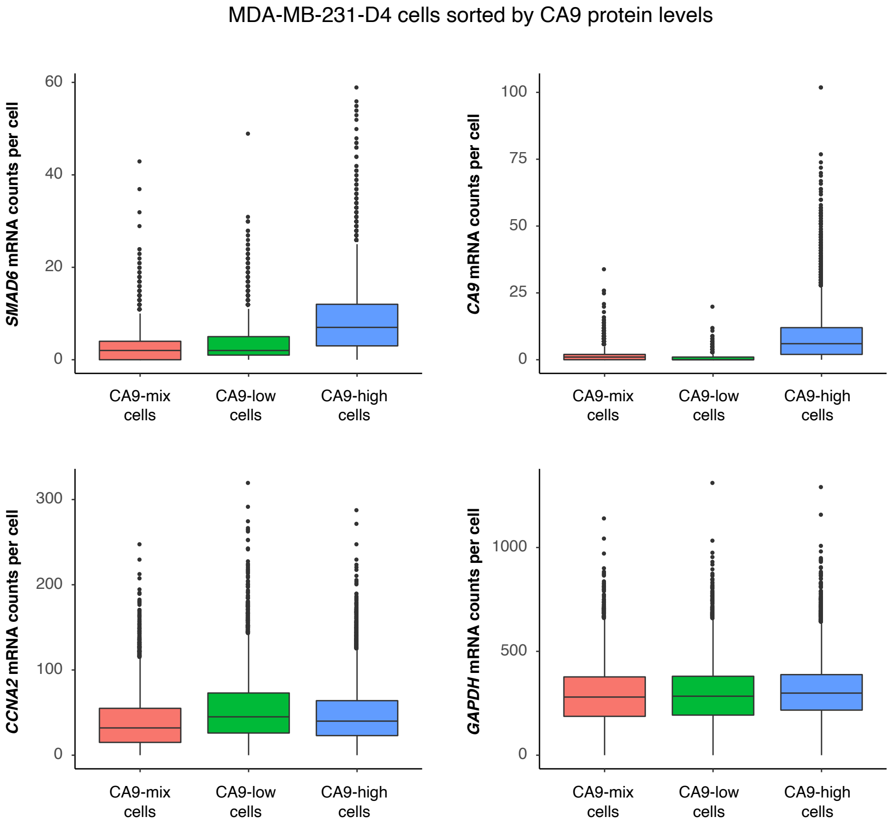


**Supplementary Figure 17. Single cell RNA FISH confirms the accuracy of sorting MDA-MB-231-D4 cells by CA9.** We sorted CA9-stained cells into high and low expressing subpopulations (as determined by antibody staining) as well as the unsorted mixed population. We then performed single cell RNA FISH for *SMAD6*, *CA9*, *CCNA2*, and *GAPDH* on these subpopulations. We found that the levels of CA9 mRNA were higher in the high subpopulation as expected, as was *SMAD6*, which was also expected based on their correlation as revealed by MemorySeq. Both housekeeping genes *CCNA2* and *GAPDH* showed minimal differences between subpopulations.

### Supplementary Figure 18

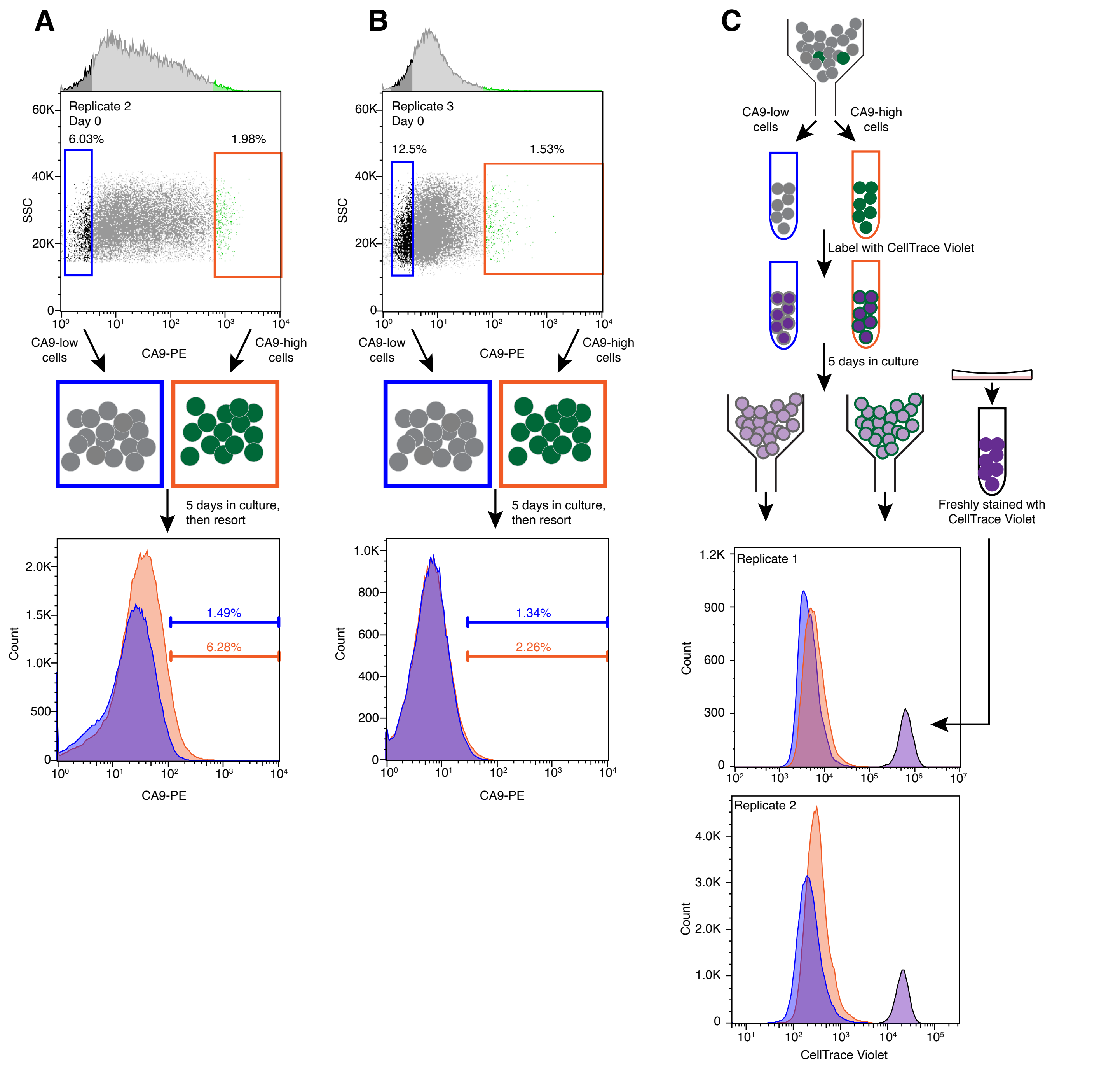


**Supplementary Figure 18. Isolated CA9-high MDA-MB-231 cells revert towards baseline CA9 levels over the course of 5 days in culture. A-B.** Biological replicates of the data presented in Figure 4C in the main text. We labeled MDA-MB-231 cells with a phycoerythrin (PE) conjugated CA9 antibody, then sorted equal numbers of cells from the top 1-2% and bottom 6-12% gates. After 5 days in culture, we re-stained the cells and measured fluorescence intensity by flow cytometry. For technical reasons, the day 5 measurement for replicate 2 was obtained on a separate flow cytometer from the day 0 sort. At day 5, the distribution of CA9-PE intensities in the CA9-high samples were shifted towards higher overall expression than the CA9-low samples, albeit with much more overlap than in the initial sort at day 0. Specifically, the top 1-1.5% gate for the CA9-low sample at day 5 contains 2-6% of the CA9-high sample. It was possible that CA9-high cells grew significantly slower than CA9-low cells, causing a decrease in frequency of the CA9-high cells over time in an inevitably impurely sorted population without those cells actually turning off. To exclude this possibility, in **C.** we monitored cell division in the CA9 sorted MDA-MB-231 by staining cells with CellTrace Violet immediately after sorting then measuring fluorescence intensity by flow after 5 days in culture. As CellTrace Violet stably labels amines in living cells, it is expected to dilute over time due to cell division. For reference, we freshly stained equal numbers of cells on day 5 to flow in parallel. As shown, both CA9-high and CA9-low sorted samples showed lower CellTrace Violet intensity than freshly stained cells consistent with multiple cell divisions (also noted by looking at clusters of cells in culture). This was true even for the cells that retained the highest levels of CA9 as measured by CA9-PE staining. While we note that the CA9-high sample retained more of the CellTrace Violet dye than the CA9-low sample, which may be due to slightly slower division or other differences in cell size or protein turnover, this difference cannot explain the change in CA9-high percentage from 100% to 2-6% over the course of 5 days.

### Supplementary Figure 19

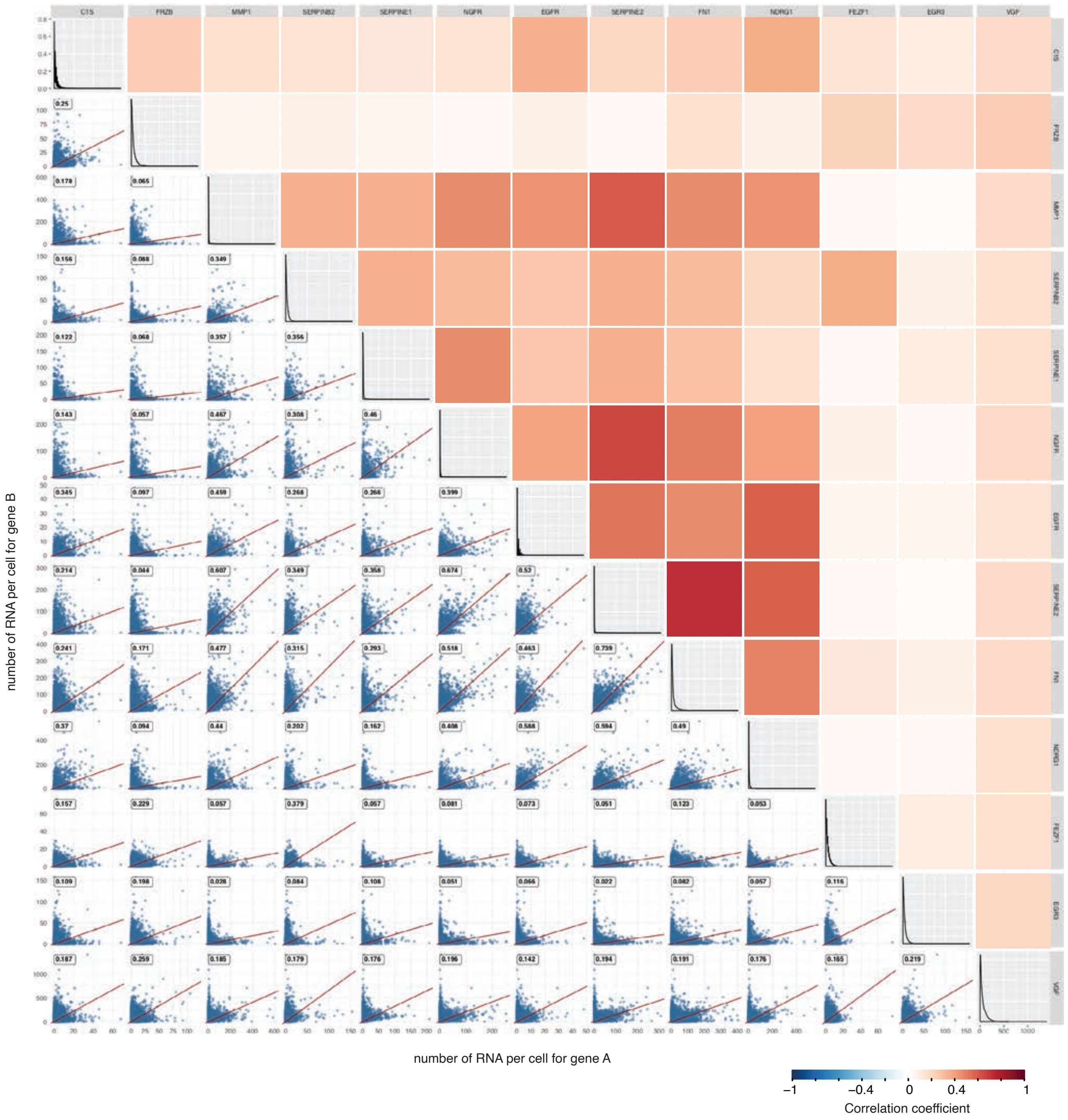


**Supplementary Figure 19. Pairwise scatterplots from the RNA FISH data shown in Fig. 5C.** Each scatterplot shows the number of mRNA per cell for each gene pair. The heatmap shows the correlation coefficient for each gene pair and is the same as the heatmap in Fig. 5C.

### Supplementary Figure 20

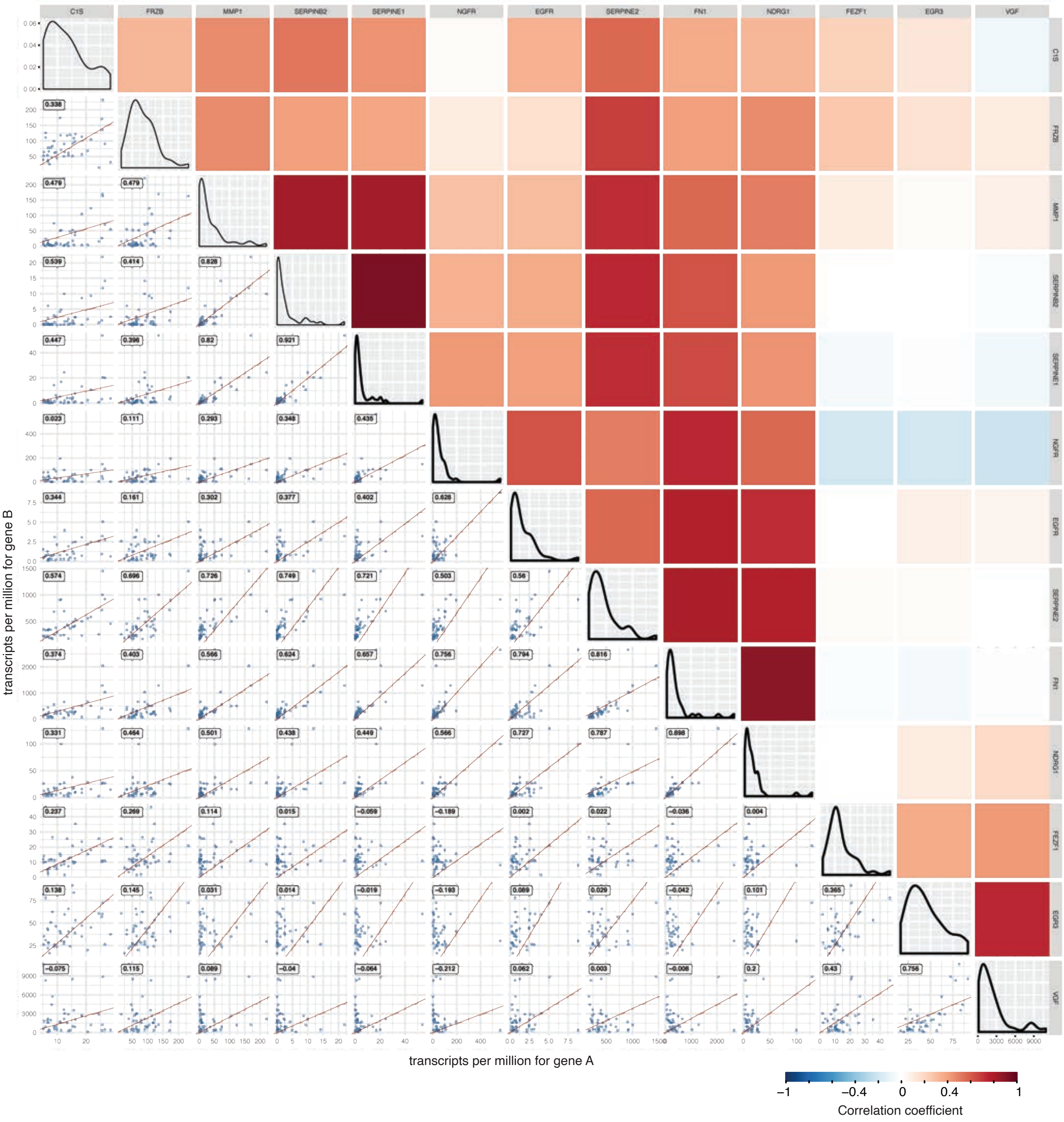


**Supplementary Figure 20. Pairwise scatterplots from the RNA-seq data shown in Fig. 5C.** Each scatterplot shows the transcripts per million of each MemorySeq clone for the gene pair. The heatmap shows the correlation coefficient for each gene pair and is the same as the heatmap in Fig. 5C

### Supplementary Figure 21

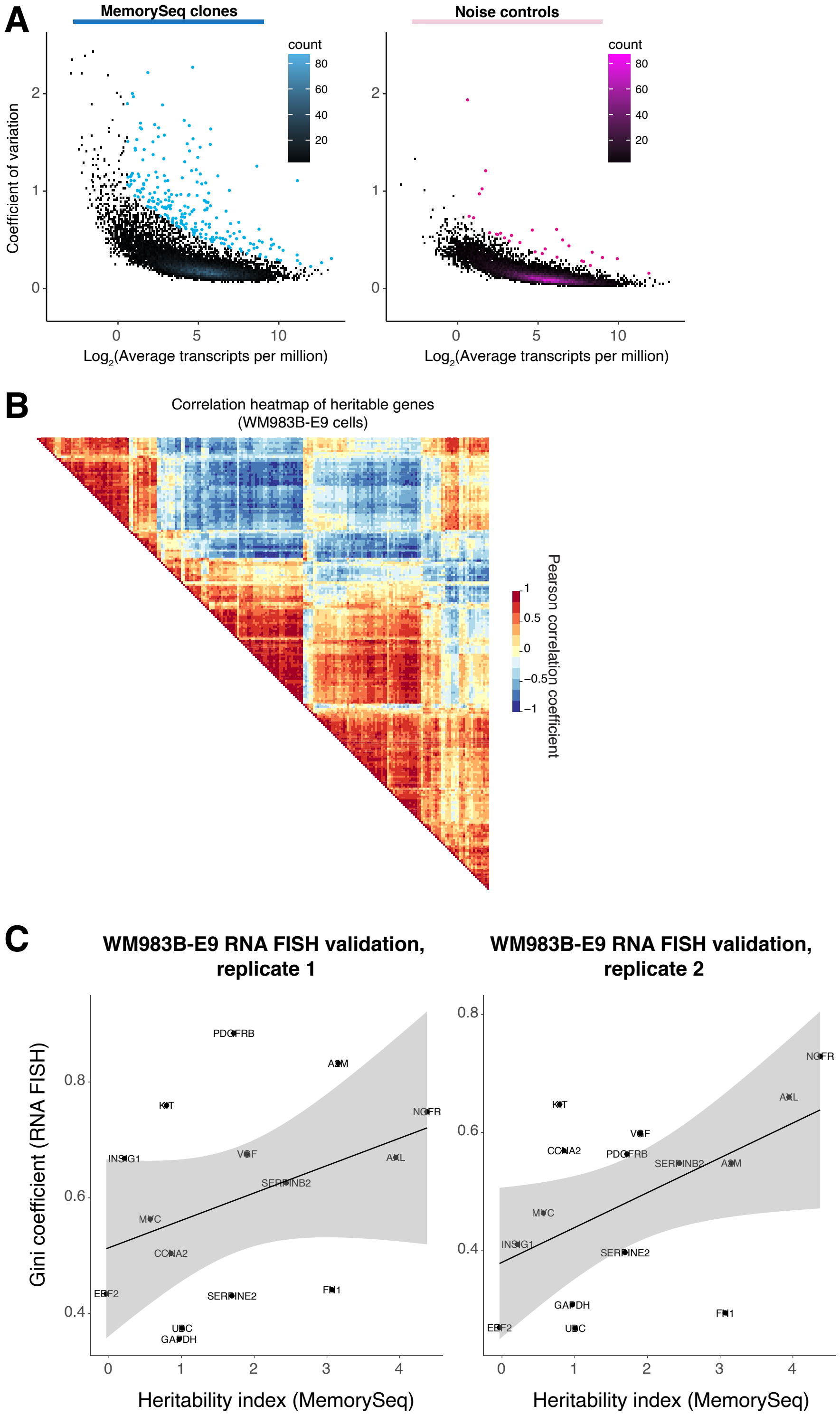


**Supplementary Figure 21. Analysis of MemorySeq data for WM983B-E9 melanoma cells. A.** We performed MemorySeq on the melanoma line WM983B-E9. Scatters of average transcript abundance from MemorySeq vs. the coefficient of variation across individual MemorySeq clones (left), with a technical noise control as described for WM989-A6 (right). **B.** Clusters of co-fluctuating genes in WM983B-E9 in genes identified by Memory- Seq as showing a high degree of heritability. **C.** We plotted heritability index from MemorySeq (skewness across individual MemorySeq clones) vs. Gini coefficient as computed from single molecule RNA FISH across two biological replicates. Overall, we found a general correspondence between the two variables, as was seen for WM983B-E9 cells, suggesting that MemorySeq also identifies genes that express in rare cells.

### Supplementary Figure 22

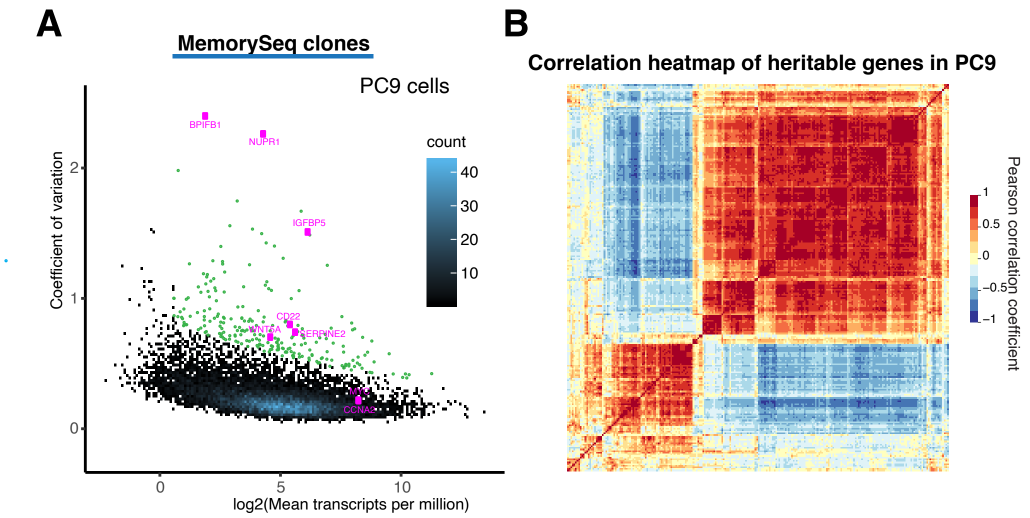


**C**

**Heritable genes in all cell lines analyzed with MemorySeq:**

*NUPR1*
*DDIT4*
*TFPI2*
*MT2A*
*CST1*
*TNC*
*FN1*
*COL6A1*

**Supplementary Figure 22. MemorySeq on PC9 reveals a set of 240 heritable genes.** We performed MemorySeq in the lung cancer cell line, PC9. **A.** Coefficient of variation versus log2 mean transcripts per million for all genes detected in PC9. Points labeled with green dots passed the threshold for being identified as a heritable gene. These genes were identified by first fitting a Poisson regression model to the data, and then selecting genes with residuals in the top 2%. The pink dots highlight a few example genes in this data set, including heritable genes (*WNT5A*, *NUPR1*, *SERPINE2*, *BPIFB1*, *CD22*, *IGFBP5*) and non-hertiable genes (*MYC*, *CCNA2*). **B.** Correlation heatmap for all heritable genes in PC9. Color represents the Pearson correlation coefficient. **C.** List of 8 heritable genes identified in all four cell lines that we analyzed with MemorySeq (WM989, WM983B, MDA-MB-231, and PC9).

### Supplementary Figure 23

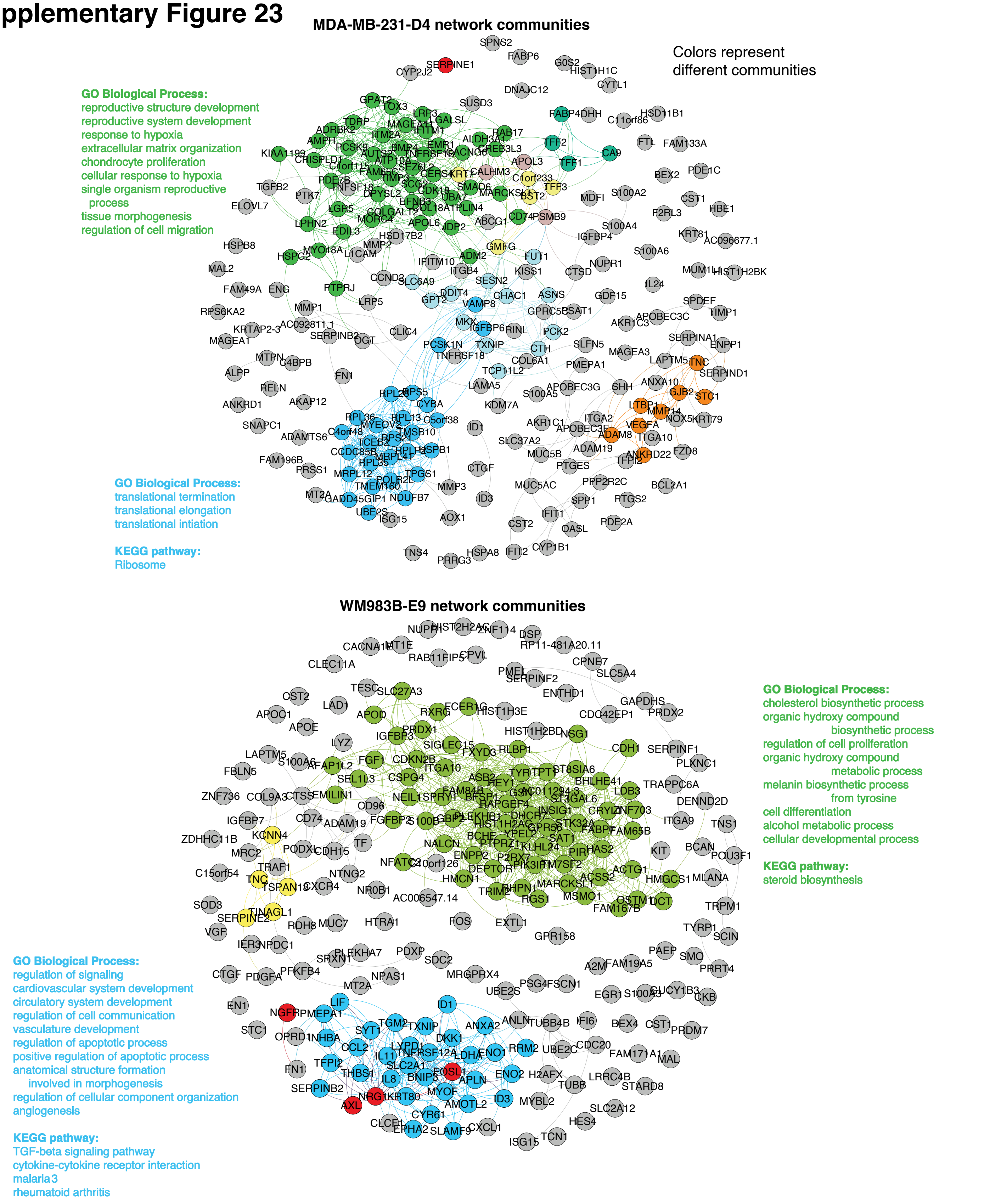


**Supplementary Figure 23. Community identification in MemorySeq hits for MDA-MB-231-D4 and WM983B.** We performed community identification on hits detected by MemorySeq (top 2% of genes, TPM of 1.5 or greater) for MDA-MB-231-D4 and WM983B-E9 cells. Cutoffs were chosen to arrive at biologically reasonable gene sets. We performed KEGG and GO biological process analysis on genes identified as belonging to particularly communities (limited to communities that had enough genes for such an analysis to be meaningful).

### Supplementary Figure 24

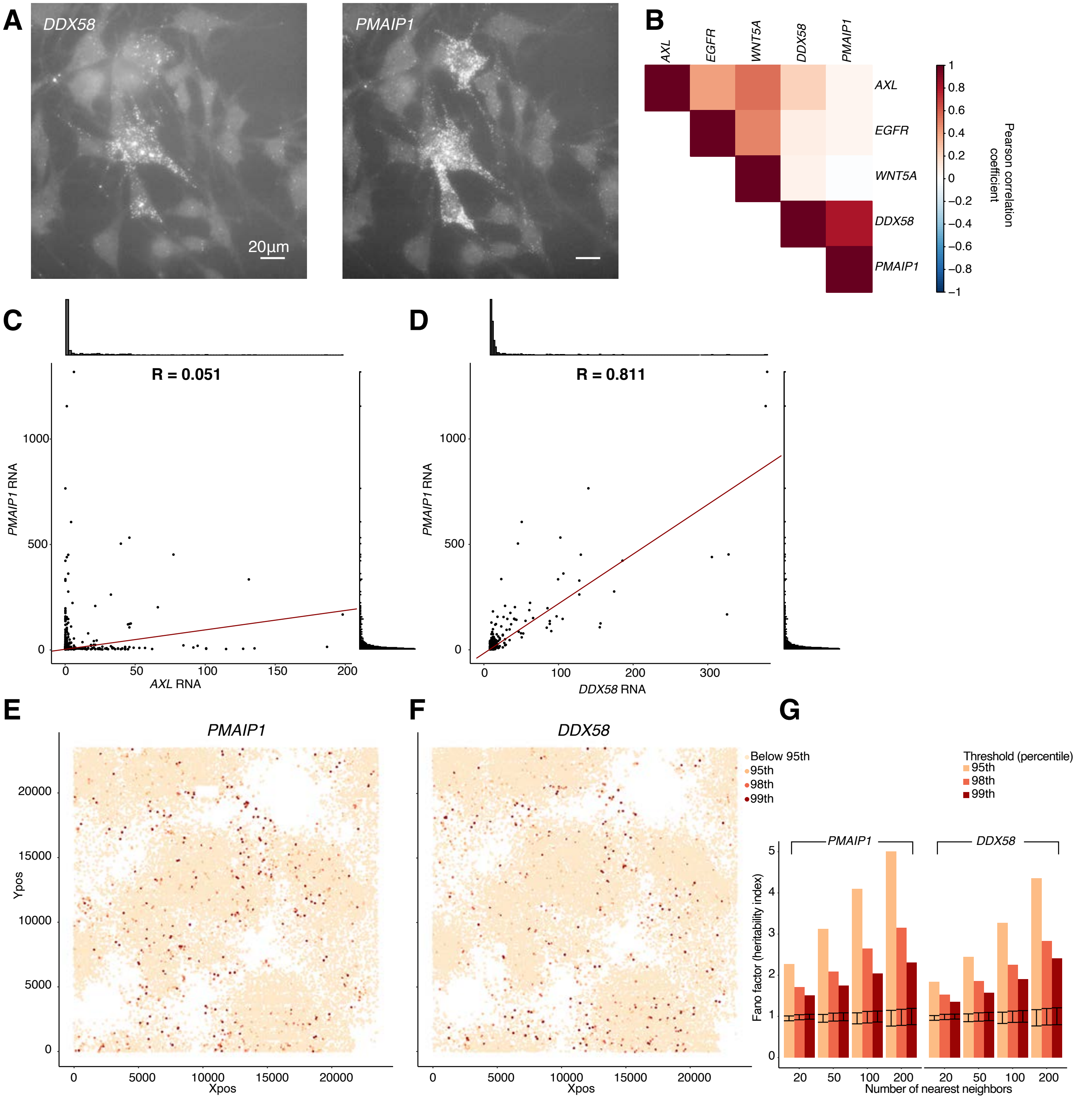


**Supplementary Figure 24. Validation of single-cell gene expression coordination and spatial clustering of *DDX58* and *PMAIP1* in WM989.** MemorySeq identified *DDX58* and *PMAIP1* as members of a correlated network of genes distinct from the network containing *AXL*, *EGFR*, *WNT5A* and other resistance markers. **A-D**. Using RNA FISH, we find that *DDX58* and *PMAIP1* expression is correlated in single-cells but both genes are far less correlated with the expression of *AXL*, *EGFR* or *WNT5A*. Shown are representative images (maximum Z-projection) of a group of cells co-expressing high levels of *DDX58* and *PMAIP1* and the measured Pearson correlation values across 362 cells. **E-G**. As described in the main text, we use spatial clustering of gene expression in single-cells as a metric for transcriptional memory. To assay spatial clustering for *DDX58* and *PMAIP1*, we sparsely plated WM989-A6-G3 melanoma then cultured the cells for 11 days before fixing and performing RNA FISH. Shown are positions of approximately 18,000 cells color coded based on their percentile of expression. For *DDX58* and *PMAIP1,* the 95th percentile corresponds to 5 and 30 RNA molecules, respectively. We quantified the significance of spatial clustering by calculating the Fano factor of the number of high expressing cells across windows of 20, 50, 100 and 200 nearest cells (see methods for further details). Error bars correspond to the 95% confidence interval of the randomly permuted dataset.

### Supplementary Figure 25

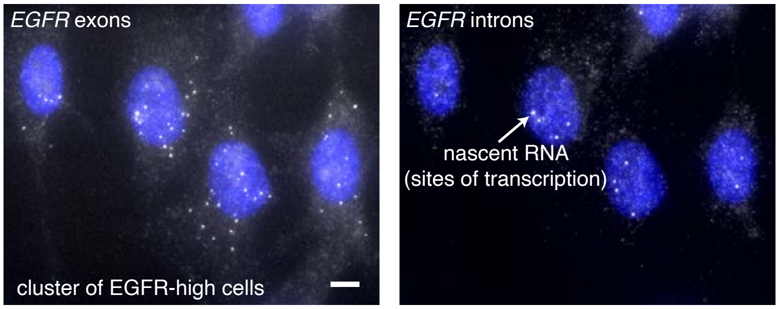


**Supplementary Figure 25. Detection of nascent transcripts show ongoing transcription in EGFR-high expressing cells.** We wondered whether EGFR-high cells (WM989-A6) stemmed from a short burst of transcription followed by dilution amongst progeny cells or from sustained transcription in this subpopulation of cells. We distinguished these possibilities by measuring nascent active transcription in single cells by targeting the intronic region of *EGFR* with single molecule RNA FISH probes in a separate color from the *EGFR* exons. We saw that in patches of EGFR-high cells, the individual cells typically also showed *EGFR* introns, showing that *EGFR* exhibited sustained transcription over multiple divisions. Scale bar is 5µm.

### Supplementary Figure 26

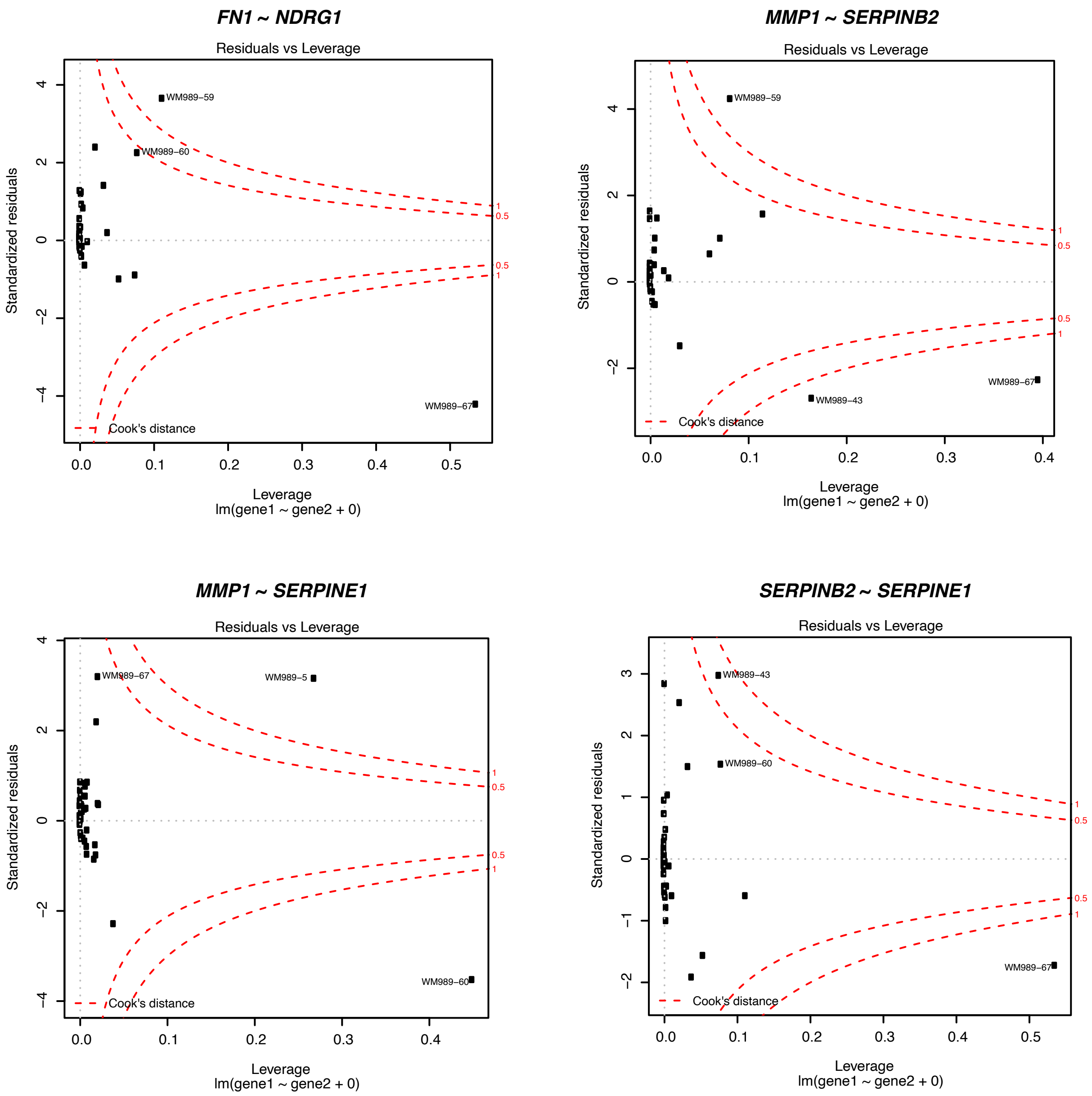


**Supplementary Figure 26. Cook's distance plots for 4 gene pairs showing the strongest correlations in the MemorySeq data.** Each plot depicts the residuals versus leverage for the regression with the gene pair. Red dotted lines delineate the cut-offs for Cook's distance > 0.5 and > 1. The MemorySeq clones with the greatest influence on the regression according to Cook's distance are labeled with the sample name.
