## Supplementary Note 1 for "Memory sequencing reveals heritable single cell gene expression programs associated with distinct cellular behaviors"

### Inferring timescale of expression heritability from MemorySeq

We model gene expression as a binary switch, where individual cells are either in an ON (high expression) or OFF (low expression) state. We assume that cells proliferate exponentially at a rate $k_x$ (i.e., a generation time of $1/k_x$), and that each MemorySeq clone begins as a single cell, ultimately growing into the final population. We make two simplifying assumptions: i) The proliferation rate of a cell is same irrespective of its ON/OFF state; and ii) The population remains in the exponential growth phase during the timespan of the experiment. Further, cells in the OFF state turn ON with rate $k_{\text{ON}}$, and revert back to the OFF state with rate $k_{\text{OFF}}$. We define $f = k_{\text{ON}}/(k_{\text{ON}} + k_{\text{OFF}})$ as the fraction of ON cells in the original population, and further assume (as indicated by our experimental data) that the ON state is rare, *i.e.*, $f \ll 1$.

Table 1: Stochastic model of cell proliferation and switching.

| Stochastic event | Population reset | Probability event occurs in small time interval $(t, t + dt)$ |
|---|---|---|
| Cell proliferation in the ON state | $\boldsymbol{x} \mapsto \boldsymbol{x} + 1$ | $k_x \boldsymbol{x} dt$ |
| Cell proliferation in the OFF state | $\boldsymbol{y} \mapsto \boldsymbol{y} + 1$ | $k_x \boldsymbol{y} dt$ |
| Cell switching from ON to OFF | $\boldsymbol{x} \mapsto \boldsymbol{x} - 1$<br>$\boldsymbol{y} \mapsto \boldsymbol{y} + 1$ | $k_{\text{OFF}} \boldsymbol{x} dt$ |
| Cell switching from OFF to ON | $\boldsymbol{x} \mapsto \boldsymbol{x} + 1$<br>$\boldsymbol{y} \mapsto \boldsymbol{y} - 1$ | $k_{\text{ON}} \boldsymbol{y} dt$ |

To perform Luria-Delbrück fluctuation analysis, we isolate a single cell from the original population, which is ON with probability $f$, and OFF with probability $1 - f$. Starting with this initial condition, we define the random variables $\boldsymbol{x}(t)$ and $\boldsymbol{y}(t)$ to be the number of cells in the ON and OFF states, respectively, at time $t$. The time evolution of integer-valued random processes $\boldsymbol{x}(t)$ and $\boldsymbol{y}(t)$ is governed by the model illustrated in Table 1. The model consists of four events that occur probabilistically with rates given in the third column, and whenever the event occurs, the cell numbers increase/decrease by one as per the reset map in the second column. The ratio $\boldsymbol{x}(t)/(\boldsymbol{x}(t) + \boldsymbol{y}(t))$ represents the fraction of ON cells at time $t$, and our goal is to quantify fluctuations in this ratio across MemorySeq clones. To do this, we derive the time evolution of the first two statistical moments of $\boldsymbol{x}(t)$ and $\boldsymbol{y}(t)$. Here we only present the main result and refer the reader to [1] for technical details on deriving moment dynamics for stochastic dynamical systems. Assuming $f \ll 1$, the coefficient of variation squared ($CV^2$) of the ratio $\boldsymbol{x}(t)/(\boldsymbol{x}(t) + \boldsymbol{y}(t))$ is given by

$$CV^2 \times f = \frac{2T_{\text{ON}} e^{T - \frac{2T}{T_{\text{ON}}}} - 2 - T_{\text{ON}}}{(2e^T - 1)(T_{\text{ON}} - 2)}, \tag{1}$$

where $T = tk_x$ is the duration of the experiment (normalized to the generation time), and $T_{\text{ON}} = k_x/k_{\text{OFF}}$ is the time spent in the ON state (normalized to the generation time). Equation (1) quantifies the noise measured in MemorySeq, and as expected, $CV^2$ is a monotonically increasing function of $T_{\text{ON}}$; *i.e.*, slower switching results in higher variation across MemorySeq clones (see figure below). It turns out that the product $CV^2 \times f$ is independent of $f$ in (1), and it is convenient to look at this product as a function of $T_{\text{ON}}$. Given *a priori* knowledge of $f$ and a measurement of the noise level $CV^2$, $T_{\text{ON}}$ can be estimated via an inverse transformation of (1). In our experiments, $f$ is typically around 1% or less, as measuring using RNA FISH, and each MemorySeq clone starts as a single cell and grows to around $10^5$ cells, for which the number of doublings $T = 17$. Using these values, the Table below estimates $T_{\text{ON}}$ for several genes based on $CV$ values across MemorySeq clones reported in Fig. 1 of the main text. To estimate the confidence intervals for $T_{\text{ON}}$, we first determine the sampling error in the measured $CV$. Assuming $T_{\text{ON}} = 6.9$ (as estimated for *EGFR* in the table below) and $T = 17$, we used the stochastic model to perform $10^3$ simulation runs, each run simulating a MemorySeq experiment with 48 clones. Our results show that 95% of $CV$s were in the range of 0.3 to 1.8. Using this range together with (1) provides a 95% confidence interval for $T_{\text{ON}} \in \{4.2, 9.6\}$ around 6.9. This analysis for the *EGFR* gene suggests a 40% uncertainty range around the estimated $T_{\text{ON}}$ values.

Table 2: Timescale of expression heritability from MemorySeq

| Gene | *WNT5A* | *NGFR* | *NDRG1* | *SERPINE2* | *EGFR* | *FOXN3* |
|---|---|---|---|---|---|---|
| MemorySeq $CV$ | 1.9 | 1.6 | 1.3 | 0.6 | 1.0 | 0.5 |
| $T_{\text{ON}}$ (# of cell generations) | 9.7 | 8.7 | 7.8 | 5.7 | 6.9 | 5.3 |

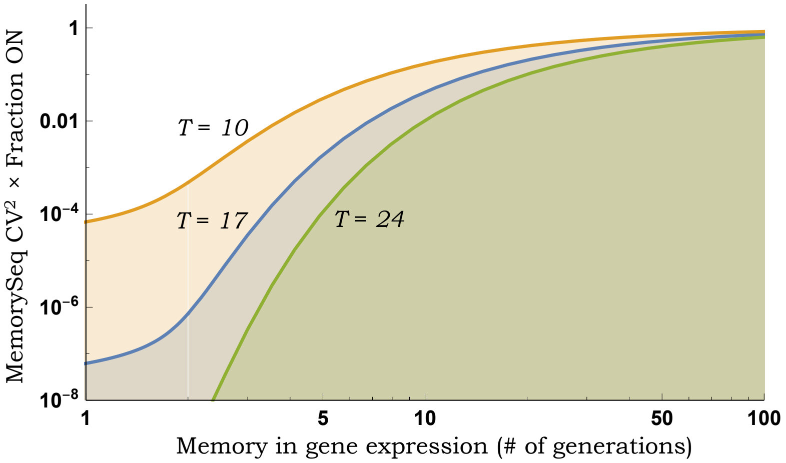


The noise $CV^2$ measured across MemorySeq clones is plotted as a function of $T_{\text{ON}}$ (average number of generations the cell remains ON) as per (1) for different durations $T$ of clonal expansion (in terms of number of cell generations). For a fixed $T_{\text{ON}}$, longer durations $T$ lead to lower noise levels.
